# Genomic and phenotypic biology of novel strains of *Dickeya zeae* isolated from pineapple and taro in Hawaii: insights into genome plasticity, pathogenicity, and virulence determinants

**DOI:** 10.1101/2021.01.28.428661

**Authors:** Gamze Boluk, Dario Arizala, Shefali Dobhal, Jingxin Zhang, John Hu, Anne M. Alvarez, Mohammad Arif

## Abstract

*Dickeya zeae*, a bacterial plant pathogen in the family Pectobacteriaceae, is responsible for a wide range of diseases on potato, maize, rice, banana, pineapple, taro and ornamentals and significantly reduces crop production; *D. zeae* causes soft rot of taro (*Colocasia esculenta*) and heart rot of pineapple (*Ananas comosus*). In this study, we used Pacific Biosciences to sequence two high quality complete genomes of novel strains of *D. zeae*: PL65 (size - 4.74997 MB; depth - 701; GC - 53.3%) and A5410 (size - 4.7792 MB; depth - 558; GC - 53.6%) isolated from economically important Hawaiian crops, taro and pineapple, respectively. Additional complete genomes of *D. zeae* representing two additional hosts (rice and banana), and other species for taxonomic comparison, were retrieved from the NCBI GenBank genome database. The genomic analyses indicated truncated type III and IV secretion systems (T3SS and T4SS) in the taro strain, which only harbors 1 and 2 genes of T3SS and T4SS, respectively, and showed high heterogeneity in the type VI secretion system. Unlike the EC1 strain, neither the PL65 nor the A5410 genome harbors the zeamine biosynthesis gene cluster, which plays a key role in bacterial virulence. The ANI and dDDH percentages between the two genomes were 94.47 and 57.00, respectively. In this study, we compared major virulence factors (plant cell wall-degrading extracellular enzymes and protease) produced by *D. zeae* strains and virulence ability on taro corms and pineapple. Both strains produced protease, pectate lyases and cellulases but no significant quantitative differences were observed (p>0.05) among the strains. All the strains produced symptoms on taro corms and pineapple leaves. Strain PL65 developed symptoms faster than the others. Our study highlights genetic constituents of pathogenicity determinants and genomic heterogeneity that will help understand the virulence mechanisms and aggressiveness of this plant pathogen.

## INTRODUCTION

*Dickeya* and *Pectobacterium,* gram negative and rod-shaped bacteria, belong to the family Pectobacteriaceae (order Enterobacterales), are devastating phytopathogens (Adeolu et al., 2016). *Dickeya* species have been listed in the top ten of most important bacterial phytopathogens because of high economic consequences (Mansfield et al., 2012). *Dickeya* currently encompasses twelve recognized species, namely *D. chrysanthemi*, *D. paradisiaca*, *D. zeae*, *D. dianthicola*, *D. dadantii* and (Samson et al., 2005), *D. solani* (van der Wolf et al., 2014), *D. aquatica* (Parkinson et al., 2014), *D. fangzhongdai* (Tian et al., 2016), *D. lacustris* (Hugouvieux-Cotte-Pattat et al., 2019), *D. undicola* (Oulghazi et al., 2019), *D. poaceiphila* (Hugouvieux-Cotte-Pattat et al., 2020), and *D. oryzae*—recently separated from *D. zeae* (Wang et al., 2020;). *Dickeya dadantii* have two subspecies, *D. dadantii* subsp. *dadantii* and *D. dadantii* subsp. *differbachieae* (Brady et al., 2012).

The *D. zeae* strains were isolated from a wide and diverse host range including pineapple, potato, maize, rice, banana, hyacinth, clivia, *Brachiaria. Chrysanthemum* and *Philodendron* (Samson et al., 2005; Sławiak et al., 2009; Toth et al. 2011; Zhang et al. 2014; Sinha et al. 1977; Li et al., 2012; Bertani et al., 2013, Pritchard et al., 2013; Hu et al., 2018). Among *Dickeya* species, *D. solani*, *D. dadantii,* and *D. zeae* usually cause serious economic crop losses, especially on potato, rice, pineapple, and banana (Hussain et al., 2008; Slawiak et al., 2009; Lin et al., 2010; Toth et al., 2011; Zhou et al., 2011; Zhang et al., 2014 and Marrero et al., 2013).

*Dickeya zeae* is diverse and complex—recently, *D. oryzae* was separated from this species (Wang et al., 2020). Bacterial strains associated with pineapple heart rot disease in Hawaii were identified as *D. zeae* as the closest match although distinguishing features appeared to warrant description as a new species (Marrero et al., 2013). Multilocus (*gap*A, *pur*A, *gyr*B, *atp*D, and *dna*A) phylogenetic analysis of pineapple strains showed high similarity with *D. oryzae* (Boluk and Arif, unpublished information).

Phytopathogens can reside on the surfaces and/or within intercellular spaces of plant leaves, without exhibiting disease symptoms (Pérombelon et al., 1992). When optimal conditions of temperature, humidity, and other factors occur, the bacteria proliferate and produce plant cell-wall-degrading enzymes (PCWDEs) leading to disease development (Hugouvieux-Cotte-Pattat et al. 1996; Pérombelon 2002). *Dickeya* species cause soft rot via coordinated production of multiple secreted enzymes, mainly PCWDEs, including pectinases, cellulases, and proteases (Hugouvieux-Cotte-Pattat et al. 1996) which constitute the primary and most essential virulence determinants (Toth et al., 2006; Davidsson et al., 2013). These PCWDEs play a significant role in bacterial pathogenesis by macerating host plant tissues, enabling host colonization and disease development (Charkowski et al., 2012; Nykyri et al., 2012; Davidsson et al., 2013). The plant cell wall is a complex of polymers (cellulose, hemicellulose, pectin, structural glycoproteins) (Pauly and Keegstra, 2016), and among these polymers, pectin is the most complex and includes both polygalacturonan and ramified regions (RGI and RGII, respectively) (Caffall and Mohnen, 2009). RGI contains a rhamnogalacturonan backbone and various lateral chains such as galactan, arabinan, and galacturonan (Caffall and Mohnen, 2009). RGII contains a short galacturonan backbone, carrying four side chains (O’Neill et al., 2004). Methyl-esterification and acetylation groups of pectin are removed by pectin methylesterases (*Pem*) and pectin acetylesterase (*Pae*) (Hugouvieux-Cotte-Pattat et al. 2014). Pectate lyases (*Pel*) cleave polygalacturonate (Hugouvieux-Cotte-Pattat et al. 2014). Unsaturated oligogalacturonates enter the periplasm using the transporters *Kdg*M and *Kdg*N (Charkowski et al., 2012). Upon entry into periplasm, these oligomers are further cleaved by polygalacturonases (*Peh*) (Hugouvieux-Cotte-Pattat et al. 2014). The small oligomers enter the cytoplasm using the transporters *Tog*T and *Tog*MNAB converted into D-galacturonate and 4-deoxy-L-to-5-hexosulose uronic acid (DKI) by the oligogalacturonate lyase (ogl) (Hugouvieux-Cotte-Pattat et al. 2014). The oligomers are catabolized into 3- phosphoglyceraldehyde by the *Kdu*ID, *Kdg*K, and *Uxa*ABC enzymes in the cytoplasm (Hugouvieux-Cotte-Pattat 2016). Additionally, D-galacturonate and 4-deoxy-L-threo-5- hexosulose uronic acid (DKI) can directly enter the cytoplasm using the transporters *Exu*T and *Kdg*T, respectively (Hugouvieux-Cotte-Pattat 2016). The rhamnogalacturonate lyase (Rhi) genes are involved in degradation of the rhamnogalacturonan I pectin-ramified regions (Hugouvieux-Cotte-Pattat et al. 2014) and resulting rhamnogalacturonan I (RGI) is cleaved by *Rhi*E that lead the oligomers and enter the cytoplasm through transporter *Rhi*T (Hugouvieux-Cotte-Pattat et al. 2014). In the cytoplasm, the enzyme *Rhi*N cleaves the unsaturated galacturonate. The periplasmic endo-galactanase (*Gan*) gene cluster is responsible for enzymes that destroy galactan chains—the galactans enter the periplasm by the transporter *Gan*L (Hugouvieux-Cotte-Pattat et al. 2014). The *Gan*A generates short oligomers that use the *Gan*FGK transport system to cross the inner membrane (Hugouvieux-Cotte-Pattat et al. 2014), and finally, the *Gan*B cleaves oligogalactan to galactose (**Figure 1**; Hugouvieux-Cotte-Pattat et al., 2001, 2014; Hugouvieux-Cotte-Pattat, 2016).

**Figure 1.**
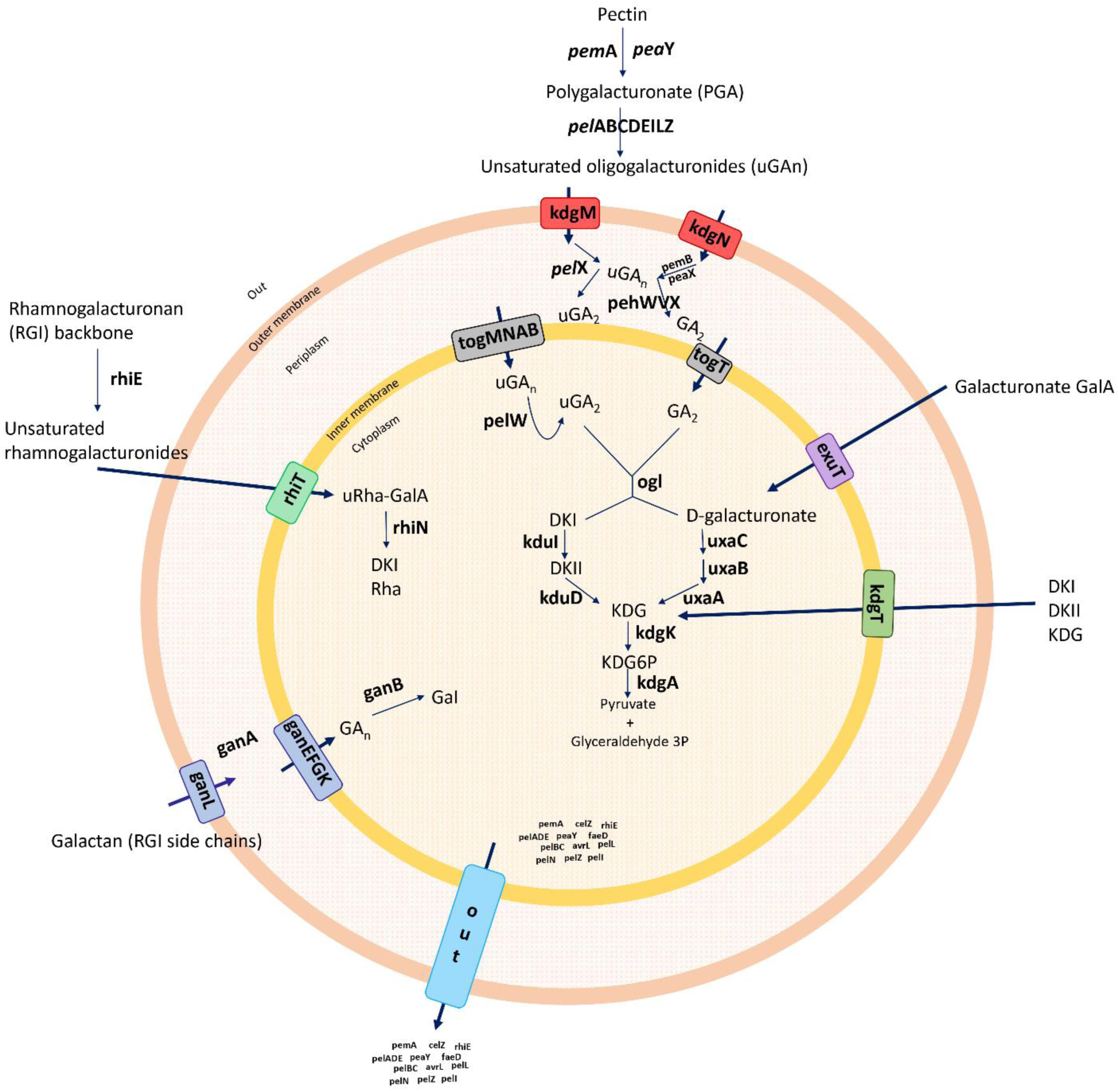
Summary of general pectin degradation pathway in soft rot bacteria.

In pectinolytic bacteria, proteases play a significant role in virulence mechanisms, and unlike the PCWDE, these enzymes are associated with the type I secretion system (T1SS) (Toth *et al*., 2006; Charkowski et al., 2012). The secretion exoenzymes from the T2SS, knows as the *out* operon, are secreted from the cytoplasm to the extracellular space (Hugouvieux-Cotte-Pattat 1996; Toth et al., 2006; Hugouvieux-Cotte-Pattat et al. 2014). Type secretion systems (T1SS – T6SS) release and modulate the transport of majority of previously described virulence components (Hugouvieux-Cotte-Pattat 1996; Toth et al., 2006). Therefore, TSS are considered the core set players that regulate the mechanism of pathogenesis in *Dickeya* (Charkowski et al., 2012). The type III secretion system (T3SS), which forms the injection machinery needed to infect plants and transport virulence proteins into the cytoplasm, is one of the major components in pathogenesis and hypersensitivity reaction (*Hrp*) (Yap et al., 2005; Alfano et al., 2004; Charkowski et al., 2012). The type IV secretion system (T4SS) referred as a conjugation system involved in bacterial DNA transfer, delivers effector proteins (virulence factors) directly to the host during infection via a cell contact-dependent way (Trokter et al., 2018). The type VI secretion systems (T6SS), possibly important for bacterial pathogenicity and host adaptation, has been associated with biofilm formation, and bacterial survival (Masum et al., 2017). In addition to TSS, bacterial pathogens form biofilms with complicated matrices including bacterial secretions that bind to the surfaces and boost the capacity of bacteria to infect the host. The biofilm development surface attachment is accelerated by functional flagella (O’Toole and Kolter 2002). Flagella and chemotaxis are essential for the establishment of successful infections (Jahn et al., 2008) Furthermore, Type IV pili are responsible for surface-associated motility (twitching motility), which allows bacteria to anchor, retract, and push forward advancing the cells (O’Toole and Kolter 2002). Cell motility, secretion, and vesicular transport are generally associated with flagellar proteins, whereas signal transduction occurs with chemotaxis proteins (Jahn et al., 2008). The pilus structures are linked to main virulence functions, namely adhesion, bacterial conjugation, surface motility, and interactions between bacteria and host cells (Craig et al., 2004; Maier and Wong, 2015). Another essential feature of biofilm development is the ability of bacteria to biosynthesize polysaccharides (Watnick et al., 2001). Exopolysaccharide synthesis plays a vital role in forming a three-dimensional architecture of biofilms (Watnick et al., 2001). The polysaccharides support multiple biological processes, such as bacterial attachment to the host, colonization, virulence and protection from plant toxins and extreme environmental conditions (Toth et al., 2006; Panda et al., 2006).

In this study, we aim first, to understand the genomic constituents of diverse *D. zeae* strains and arsenals involved in pathogenicity and host adaptation through comparative genomic analyses; and secondly, to understand genomic and phenotypic biology of the two novel strains, PL65 and A5410, isolated in Hawaii from taro and pineapple, respectively.

## 2. MATERIALS AND METHODS

### 2.1. Bacterial strains and genomic DNA extraction

Two novel strains (A5410 and PL65) of *Dickeya* species were isolated from two different hosts and both isolated from Hawaii. Strain A5410 was isolated in August 2007 from pineapple fruit (*Ananas comosus*) showing symptoms of pineapple heart rot. Pineapples, planted with imported suckers from Philippines, exhibited heart rot symptoms in Hawaii (Sueno et al. 2013). Strain PL65 was isolated in 2018 from a taro corm (*Colocasia esculenta*), showing soft rot symptoms. These strains are available at the Pacific Bacterial Collection, University of Hawaii at Manoa.

Bacteria were streaked on dextrose peptone agar (DPA: peptone 10 g/l, dextrose 5 g/l, and agar 17 g/l) (modified from Norman and Alvarez, 1989) and incubated at 28°C for 24 h. A single colony was streaked onto DPA and incubated at 28°C for 24 h.

A half-loopful overnight grown bacterial culture was used to extract the genomic DNA using the QIAGEN Genomic-tip 100/G (Qiagen, Valencia, CA) according to the manufacturer’s instructions. Quantification and quality control of the DNA was performed using a Nanodrop spectrophotometer, and a Qubit 4 fluorometer (Thermo Fisher Scientific, Life Technologies, Carlsbad, CA).

### 2.2. Whole-genome sequencing and annotation

The whole-genome sequencing of both strains was performed at the Washington State University using a PacBio RS II (Pacific Biosciences of California, Inc., Menlo Park, CA) with a Single-Molecule Real-Time (SMRT). The libraries were prepared with a 20 kb insert-size and sequenced using C4 sequencing chemistry and P6 polymerase. The sequencing reads were trimmed based on quality and length to generate highly accurate long reads and further assembled using the Hierarchical Genome Assembly Process HGAP v4 (Pacific Biosciences, SMRT Analysis Software v2.3.0). The assembled genomes were annotated using three different pipelines: the NCBI Prokaryotic Genome Annotation Pipeline (PGAP) (Tatusova et al., 2016), the Integrated Microbial Genomes pipeline version 4.10.5 from the Joint Genome Institute (IMG-JGI; Huntemann et al., 2015), and the Rapid Annotation System Technology (RAST) server (Brettin et al., 2015).

### 2.3. Comparative genomics and phylogenomic analyses of *Dickeya* species

Thirteen complete and high-quality draft genomes, including *Dickeya aquatica* 174/2, *D. chrysanthemi* Ech1591, *D. dadantii* 3937, *D. dianthicola* ME23, *D. fangzhongdai* PA1, *D. lacustris* S29, *D. paradisiaca* Ech703, *D. solani* IPO 2222, *D. undicola* FVG10-MFV-A16, *D. zeae* EC1, Ech586 and MS2 and *Pectobacterium atrosepticum* 36A, were retrieved from the NCBI GenBank genome database on February 02, 2020 (Table 1).

**Table 1.** Genomes selected for comparative and phylogenomic analyses.

| Species | Strain Name | NCBI Accession Number | Location | Host/Source | Isolation Year | Replicons (Assembly Level) | Genome Size (Mb) | GC% |
| --- | --- | --- | --- | --- | --- | --- | --- | --- |
| *Dickeya zea* | EC1* | NZ_CP006929 | China | *Oryza sativa* | - | 1 contig | 4.53 | 53.4 |
| *Dickeya* sp. | A5410 | NZ_CP040817 | USA | *Ananas comosus* | 2007 | 1 contig | 4.78 | 53.5 |
| *Dickeya* sp. | PL65 | NZ_CP040816 | USA | *Colocasia esculenta* | 2018 | 1 contig | 4.75 | 53.6 |
| *D. zea* | Ech586 | NC_013592 | USA Florida | *Philodendron* Schott | - | 1 contig | 4.82 | 53.6 |
| *D. zea* | MS2 | NZ_CP025799 | China | *Musa* sp. | 2014 | 1 contig | 4.74 | 53.4 |
| *Dickeya aquatica* | 174/2<sup>T</sup> | NZ_LT615367 | UK | River water | 2012 | 1 contig | 4.5 | 53.6 |
| *Dickeya chrysanthemi* | Ech1591 | NC_012912 | - | *Zea mays* | - | 1 contig | 4.81 | 54.5 |
| *Dickeya dadantii* | 3937 | NC_014500 | France | *Saintpaulia ionantha* | 1977 | 1 contig | 4.92 | 56.3 |
| *Dickeya dianthicola* | ME23 | NZ_CP031560 | USA Maine | *S. tuberosum* | 2016 | 1 contig | 4.91 | 55.7 |
| *Dickeya fangzhongdai* | PA1 | NZ_CP020872 | China | *Phalaenopsis* sp. | 2011 | 1 contig | 4.98 | 56.9 |
| *Dickeya lacustris* | S29<sup>T</sup> | NZ_QNUT01 | France | River water | 2017 | 118 contigs | 4.31 | 53.1 |
| *Dickeya paradisiaca* | Ech703 | NC_012880 | Australia | *S. tuberosum* | - | 1 contig | 4.68 | 55 |
| *Dickeya solani* | IPO 2222<sup>T</sup> | NZ_CP015137 | Netherland | *S. tuberosum* | 2007 | 1 contig | 4.92 | 56.2 |
| *Dickeya undicola* | FVG10-MFV-A16 | NZ_RJLS00 | France | Fresh water | 2017 | 202 contigs | 4.54 | 54.5 |
| *Pectobacterium atrosepticum* | 36A | NZ_CP024956 | Belarus | *S. tuberosum* | 1978 | 1 contig | 4.97 | 51.1 |

Type strains are marked with “<sup>T</sup>” after the strain name.
The data that is not available are marked with “-”
\*It was *Dickeya zea*, but recently, it has been reclassified as *D. oryzae*

Pairwise comparison of A5410 and PL65 genomes with the thirteen genomes was performed using average nucleotide identity (ANI) based on the Nucleotide MUMmer algorithm (ANIm) in JSpecies Web Server (Richter et al. 2016). The digital DNA-DNA hybridization (dDDH) was calculated using the Genome-to-Genome Distance Calculator (GGDC) (http://ggdc.dsmz.de/ggdc.php#) version 2.1 with the recommended formula two and BLAST+ alignment criteria. The ANI and dDDH data were compiled in a single matrix and visualized as a color-coded heatmap using DISPLAYR (https://www.displayr.com/). The cut-off values of 95- 96% (Richter et al., 2009; Goris et al., 2007; Kim et al., 2014; Chun et al., 2018; Wang et al. 2020) and 70% (Wayne 1987; Goris et al., 2007) were assigned as a species delineation framework for ANI and dDDH, respectively.

Blast matrix, codon usage, amino acid usage, and pan-core analyses across *Dickeya* strains were analyzed using the CMG-Biotools pipeline (Vesth et al. 2013). The percentage of shared proteins among the *Dickeya* strains was computed based on 50/50 Blast analysis (50% identity match and 50% length identity). The generated BLAST matrix plot was visualized as a color scale heatmap showing the numerical homology percentages across all compared proteomes. Besides, a clustering analysis according to codon and amino acid usage data was determined for genomes that displayed DDH, ANI, ANIm, and TETRA values below to the cut-off parameter for species delineation compared with the reference strain of affiliated *Dickeya* species. The codon and amino acid usage were calculated using BioPerl modules (Stajich et al., 2002) and visualized as heatmaps using R as implemented in the CMG-Biotools (Vesth et al., 2013).

The pan-core genome plot, tree analyses, and predicted proteome comparisons were performed with 14 *Dickeya* species genomes using CMG-Biotools (Vesth et al., 2013). Pairwise pan- and core-genomes were calculated for all genome combinations as mentioned above using the BLAST algorithm (Basic Local Alignment Search Tool) (Altschul et al., 1990) with a 50% of cut-off values for either query cover or identity percentage parameters. The core- and pan-genome plots were visualized in the pancoreplot program using CMG-biotools (Vesth et al., 2013).

The phylogenetic relationship was performed based on multilocus sequence analyses using 86 virulence-related genes (cell wall-degrading enzyme genes) (Table S1). The corresponding gene sequences of fourteen *Dickeya* species and one *Pectobacterium* species (used as an out-group) were retrieved from the NCBI GenBank genome database (Table 1). The 86 concatenated gene sequences were aligned using progressive Mauve plugin in GENEIOUS Prime v 2020.0.4. The concatenated alignment data was used to generate the Neighbor-Joining phylogenetic tree using CLC Genomics Workbench 20 (Qiagen).

### 2.4. Genome comparisons of *Dickeya zeae* species complex

The complete genomes of both novel *Dickeya* sp. strains (A5410 and PL64) were compared with three other complete genomes of *D. zeae* (EC1, Ech586, and MS2). Recently, *D. zeae* strain EC1 was reclassified as *D. oryzae* (Wang et al., 2020), a novel species within genus *Dickeya*. However, we included the genome of EC1 in our analyses because of the close relationships with *D. zeae* strains. Complete genomes of EC1, Ech586, and MS2 were retrieved from the NCBI genome database. The basic genomic profile features of two novel strains were taken from the NCBI GenBank database and the Bioinformatic Resource Center PATRIC Web server (Wattam et al., 2014; Wattam et al., 2017) (Table S1). Additionally, the genome atlases were constructed to illustrate different structural components present in the DNA sequences such as GC skew, stacking energy, intrinsic curvature, position preference, global direct and indirect repeats. The previous parameters were visualized and drawn as a circle plot using the GeneWiz program (Hallin et al., 2009), which were outputted in the workbench CMG-Biotools by using the script atlas_createConfig (Vesth et al., 2013). Genomic islands (GI) were predicted using the IslandViewer 4 webserver (Bertelli et al., 2017) for both new strains. IslandViewer 4 (IslandPath-DIMOB (Bertelli et al., 2018), SIGI-HMM, IslandPick (Langille et al., 2008), and Islander (Hudson et al., 2015)) were used in order to generate an interactive visualization of genomic islands (GI) by using.

Multiple genome alignment of all five genomes was conducted using the progressiveMauve 2.3.1 (Darling et al. 2004). The set of common (core genome) and unique genes within the *Dickeya* genus were identified using an all-against-all comparison determined with OrthoMCL pipeline using the BLASTP algorithm all-against-all genomes comparison included in this study (Li et al., 2003). The orthologous gene clustering analyses were implemented with default settings. OrthoMCL clustering analyses were performed with following parameters: P-value Cut-off = 1 × 10^−5^; Identity Cut-off = 90%; Percent Match Cut-off = 80 (Li et al., 2003).

The clusters involved in various virulence and pathogenicity functions (such as the plant cell wall degrading enzymes (PCWDE), type secretion systems (I-VI), synthesis of polysaccharides (enterobacterial common antigen, capsular polysaccharide, lipopolysaccharides, exopolysaccharides, and O-antigen), bacterial attachment operons (type IV pili), flagella and chemotaxis) were screened and compared among the *D. zeae* complex using the Proteome Comparison tool of Pathosystems Resource Integration Center (PATRIC) web server (Wattam et al., 2017). The synteny and different rearrangements between the main pathogenicity genomic clusters were visualized as linear arrows generated using Easyfig v2.2.3 (Sullivan et al., 2011).

The secondary metabolites biosynthetic related gene clusters were predicted using antiSMASH 4.0 (Blin et al., 2017). The Clustered Regularly Interspaced Short Palindromic Repeats (CRISPRs) arrays and the type of CRISPR-associated proteins (Cas) systems were predicted using CRISPRCasFinder (Couvin et al., 2018).

### 2.5. Phenotypic comparisons of novel strains

A type strain, NCPPB 2538 (A5422=CFBP 2052), of *D. zeae* along with A5410 and PL65 were phenotypically characterized. The complete genome of the type strain (NCPPB 2538) was not available in public databases, and therefore, we did not include this genome in the phenotypic analyses. The selected three strains were characterized for production of extracellular enzymes, swimming and swarming ability, polysaccharides synthesis, biofilm formation, and plant infection ability following different protocols as mentioned below. Initially, the growth curve pattern was evaluated for all three strains.

#### 2.5.1. Pel, Cel, and Prt enzyme activity assays

The protocol reported by Chatterjee et al., 1995 was followed for the enzyme activity assays. Briefly, media composition (per liter) pectate lyase (*Pel*) assay medium—10 g polygalacturonic acid (PGA), 10 g yeast extract, 0.38 µmol CaCl_2_ and 100 mmol Tris-HCl, pH 8.5, 8 g agarose and 2 g sodium azide; cellulase (*Cel*) assay medium—1 g carboxymethyl cellulose and 25 mM sodium phosphate, pH 7.0, 8 g agarose and 2 g sodium azide; protease (*prt*) assay medium—30 g gelatin, 4 g nutrient 8 g agar and 2 g sodium azide. The media plates were prepared by pouring, and 3 mm diameter wells were punched after solidification, and bottoms were sealed with molten agarose. The bacterial suspension (50 µl of overnight culture; OD_600_∼2.2) was applied to each well, and plates were incubated at 28°C. After 10h, Pel assay plates were flooded with 4 N HCl, and Cel assay plates were flooded with 2% Congo Red solution for 10 min and washed for 5 min with 5 M NaCl. Haloes around the wells indicated protease activity within 24 h without any further treatment. The diameter of a clear halo around the colonies was measured. Each treatment was carried out in triplicates, and all assays were repeated three times.

#### 2.5.2. Motility assay

Swimming and swarming motility assays were performed in semisolid media plates as described by Chen et al., 2016. The single-pure bacterial colony was inoculated in the middle of the plates (swimming medium per liter contained 10 g tryptone, 5 g NaCl, and 3 g agar and supplemented with 0.05% (w/v) tetrazolium chloride while swarming medium per liter contained 10 g tryptone, 5 g NaCl, and 4 g agar and supplemented with 0.05% (w/v) tetrazolium chloride) using a toothpick. Plates were incubated at 28 °C for 24 hours, and the diameter of the bacterial growth was measured. Each treatment was carried out in triplicates, and assays were repeated three times.

#### 2.5.3. Exopolysaccharides production assay

The exopolysaccharide production assay was performed in solid medium plates following the procedure described by Narváez-Barragána et al., 2020. A single-pure bacterial colony was inoculated in the middle of solidSOBG plates (solidSOBG medium each liter contained 20 g tryptone, 5 g yeast extract, 10 mM NaCl, 2.5 mM KCl, 10 mM MgSO4, 15 g agar supplemented with 2% glycerol) using a toothpick. Plates were incubated for 3 days at 28 °C. The width of the line was measured to calculate EPS production. Each treatment was carried out in triplicate, and assays were repeated three times.

#### 2.5.4. Biofilm formation and quantification assays

The biofilm assay was performed in SOBG (Super Optimal Broth with 2% glycerol) as described by Chen et al. 2016 with minor modifications. The overnight grown bacterial culture was diluted in 1:100 in SOBG broth (SOBG contained the same component of solid SOBG medium without agar). The 96 well microliter plates containing 100 μl of diluted culture in each well was incubated at 28°C, shaking at 200 rpm for 18 h. Then, bacterial cultures were removed and added 125 μl of 0.1% crystal violet (w/v). After 15 min staining at room temperature, the dye was washed three times with water. Stained wells were decolorized with 200 μl of 95% ethanol, after drying, attached bacterial cells were quantified. The quantity of crystal violet was determined by measuring the absorbance at 600 nm using a BioTek Epoch Microplate Spectrophotometer (Vermont, USA).

#### 2.5.5. Pathogenicity assay

Pathogenicity assays were performed on taro corms and pineapple leaves to compare the virulence using both novel strains and a type strain (NCPPB 2538 (A5422=CFBP 2052)) as a reference. Plant materials were washed carefully under running water, and surface sterilized with 0.6% sodium hypochlorite for 10 minutes. Later, plant materials were cleaned and left in water for 10 min and dried inside the laminar flow. Dried plant materials were infected using 50 μl of overnight grown culture (OD_600_∼2.2); bacteria were grown in NBG (Nutrient broth + 0.4 w/v % glucose). The infected plant materials were incubated inside a moist chamber at 25 °C for 48 h. The controlled plant materials were inoculated with NBG and incubated as described above. After 48 h incubation, macerated taro tissues were removed and weighted (g) without drying, to determine virulence levels of the strains. In other pathogenicity assays, at least two pineapple leaves were inoculated with ¼ loop of overnight grown culture. Inoculated leaves were incubated at room temperature for 48 h. Each treatment was repeated three times, also each experiment was repeated three times.

### 2.6. Data Analyses

Each experiment was performed in three replicates. Analysis of variance was calculated using the IBM SPSS Statistics V25 (IBM Corp. Released 2017. Version 25.0. Armonk, NY) one-way analysis of variance (P < 0.05). Mean value differences were calculated by the Tukey test.

## 3. RESULTS

### 3.1. Comparative genomics and phylogenomic analyses within *Dickey*a Genus

Average nucleotide identity (ANI) and *in-silico* DNA-DNA hybridization (dDDH) analyses showed that the five *D. zeae* genomes were distinct from the other well-characterized species (**Figure S1**). High genome dissimilarity was observed among the *D. zeae* species complex with a dDDH value of 56-68.20 %, which is lower than the recommended cut-off value (70%) (Wayne et al., 1987; Goris et al., 2007). Additionally, the ANI value within the *D. zeae* species was 94.33- 96.27%, which overlaps the threshold value (ANI 95–96%) (Richter et al., 2009; Goris et al., 2007; Kim et al., 2014; Chun et al., 2018; Wang et al. 2020). The dDDH and ANI analyses indicated that the strains of *D. zeae* complex shared highest DNA homology with *D. chrysanthemi* Ech1591 (dDDH 30.40-30.9%; ANI 87.24-87.59%). The ANI and the dDNA-DNA hybridization values between the *Dickeya* and *Pectobacterium* genus were 83.68-84.48% and 20.70-21.60%, respectively (**Figure S1**). The dDDH phylogenetic tree was inferred with FastME 2.1.6.1 from GBDP distances calculated from genome sequences (Meier-Kolthoff et al 2013). The ANI phylogenetic tree was generated for the *Dickeya* species strains based on whole-genome alignment using the neighbor-joining method. The Jukes-Cantor model was used for analysis, and the tree was built based on 1,000 bootstraps—CLC Workbench 20 was used for analyses. The *D. zeae* complex was clustered, and *D. chrysanthemi* was the closest related species in the phylogenetic tree (**Figure S2A-B**). Interestingly, while *P. atrosepticum* and *D. paradisiaca* grouped together according to ANI-NJ phylogenetic analyses, in the dDDH phylogenetic analysis *P. atrosepticum* was an outgroup strain (**Figure S2A-B**).

The phylogenetic analyses based on the concatenated sequences of eighty-six virulence-associated genes (Table S1) were used to evaluate the evolutionary relationships between *D. zeae* complex and other members of *Dickeya*. The phylogenetic tree revealed two main separate clades: the first (**Figure 2A**) grouped all *D. zeae* complex as well as *D. dianthicola, D. undicola, D. fangzhongdai, D. solani, D. dadantii,* and *D. chrysanthemi*; the second clade consisted of *D. paradisiaca, D. lacustris, D. aquatica* and *P. atrosepticum*.

**Figure 2.**
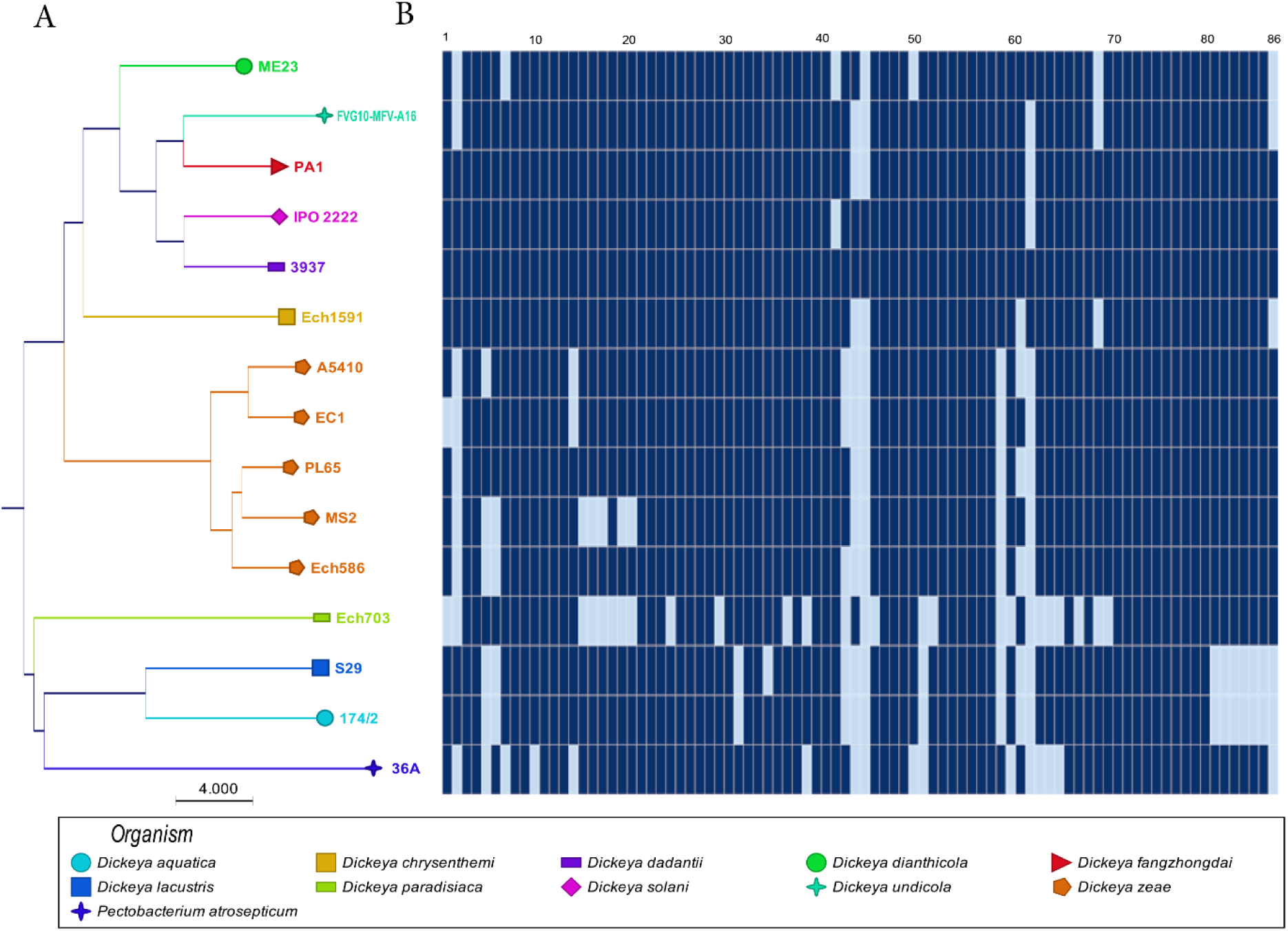
(**A**) Concatenated Neighbor-Joining phylogenetic dendrogram based on 86 plant cell wall degrading enzymes in *Dickeya* sp. The Neighbor-Joining method was applied to determine the evolutionary distances, with each node being supported by a bootstrap of 1,000 replicates to assess reliability. (**B**) Heat map based on present and absent 86 plant cell wall degrading genes in the genomes. Dark blue-shows present genes, and light blue-shows absent genes. In Figure 2B, number 1-86 are described in Table S1.

### 3.2. The BLAST matrix analysis in *Dickeya* genus

The BLAST matrix heatmap was generated to determine the similarity in each of the conserved protein families present within the fourteen *Dickeya* species. The pairwise comparison of total protein-coding genes among the fourteen *Dickeya* genomes ranged 51.1% to 77.8% shared proteins, with the lowest value representing the pairwise identity between *D. aquatica* and *D. paradisiaca* and the highest between PL65 and Ech586 *D. zeae* genomes (**Figure 3**). The number of proteins and protein families used to compare all proteomes were the lowest for the genome EC1 of *D. zeae* with 3,887 proteins and 3,711 families (**Figure 3**). The proteome comparison displayed that the average protein family similarity among *D. zeae* genomes ranged 71.5 % to 77.8%. In comparison, the intra-proteome homology among protein families within each genome is less than 3.5% (**Figure 3**). These results indicated that *D. zeae* is genetically distinct from other species. The blast matrix results are concordant with the ANI and dDDH analyses. The BLAST matrix results also demonstrated that *D. zeae* proteomes are diverse, with an average of 74% sequence identity, one exception was the sequence identity of 77.8% between the Ech586 and PL65 genomes; this could be because these strains were isolated from hosts in the Araceae family. (*Colocasia esculenta*-PL65 and *Philodendron* Schott-Ech586). Overall, these results suggest there was a significant diversity among strains in the *D. zeae* species complex.

**Figure 3.**
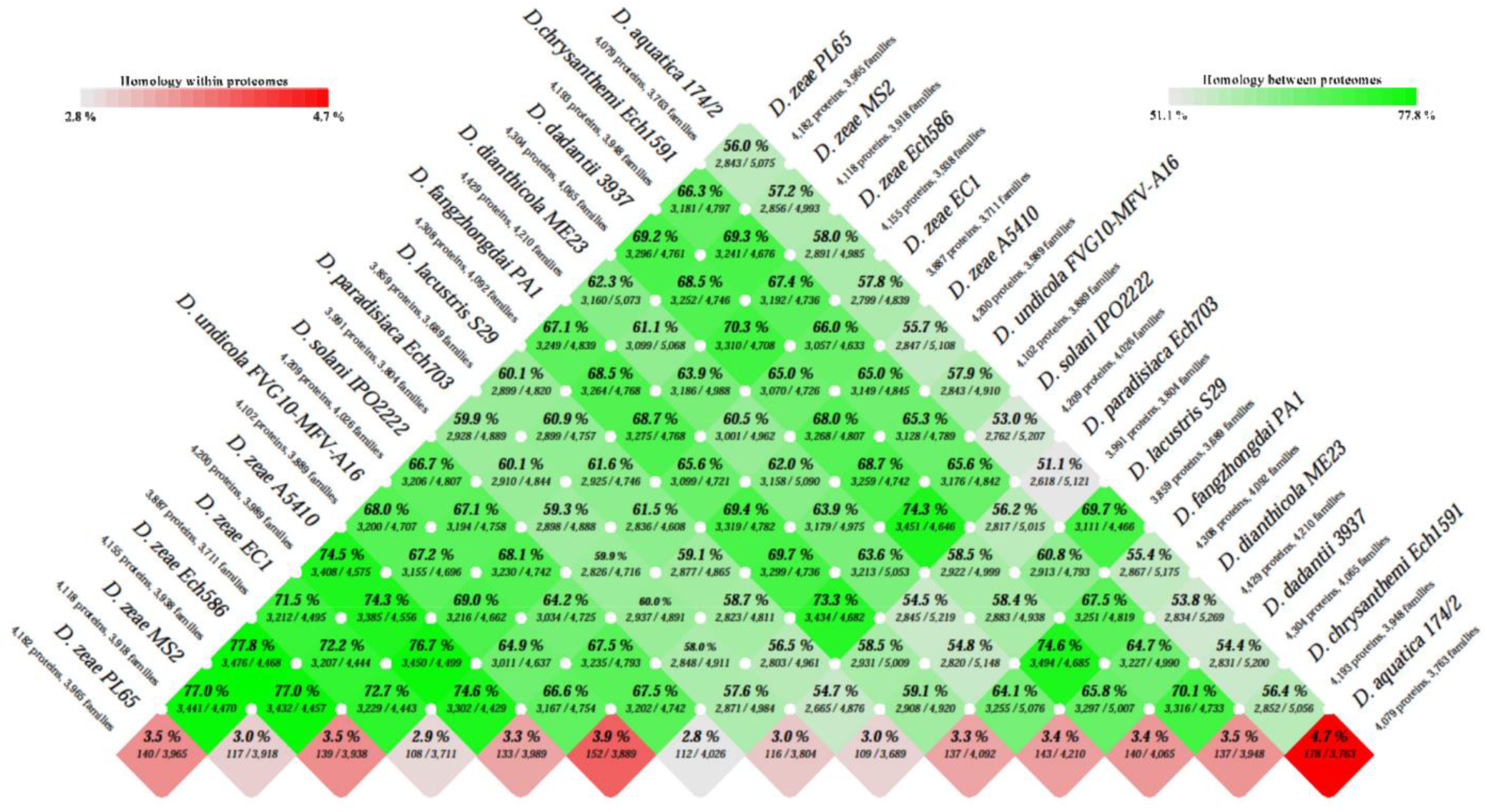
BLAST matrix between and within total proteomes of *Dickeya* genus. Pairwise protein comparison was performed using BLAST. All protein coding sequences were compared with each other across the genomes. The BLAST hit was considered significant when 50% of the alignment showed identical matches and if coverage of the alignment was at least 50%. The color scale intensity from dark green to light green highlights the degree of homology between the proteomes whereas the color scale from dark red to light red shows the homologous hits within the proteome itself.

### 3.3. Genomics evolution of *Dickeya* species based on analysis of codon and amino acid usage

Same amino acid can be generated by more than one codon, and these codons are referred as synonymous codons (Uddin, 2017). The codon usage shows the variation among the genomes and species. Hence, the codon usage pattern establishes a unique characteristic of each species (Uddin, 2017). In this concept, we analyzed and contrasted the bias in the third codon position, and amino acid frequencies for the fourteen *Dickeya* species. The corresponding analysis consisted of quantifying the fraction of each codon and amino acid count of the total number of codons and amino acids. The percentage of codon and amino acid profiles among the species was calculated and visualized in heatmaps (**Figure 4A** **and 4B**). Codon usage heatmap displayed a yellow color intensity for the usage of GC-rich codons like GCG, CGC, CTG, CAG, CGG, CCG, GGC, and GCC heatmap (**Figure 4A**). The amino acid usage heatmap displayed that amino acids like Alanine (A), Arginine (R), Leucine (L), and Serine (S) (indicated as pink color scales) were used in higher frequency in the *Dickeya* species (**Figure 4B**).

**Figure 4.**
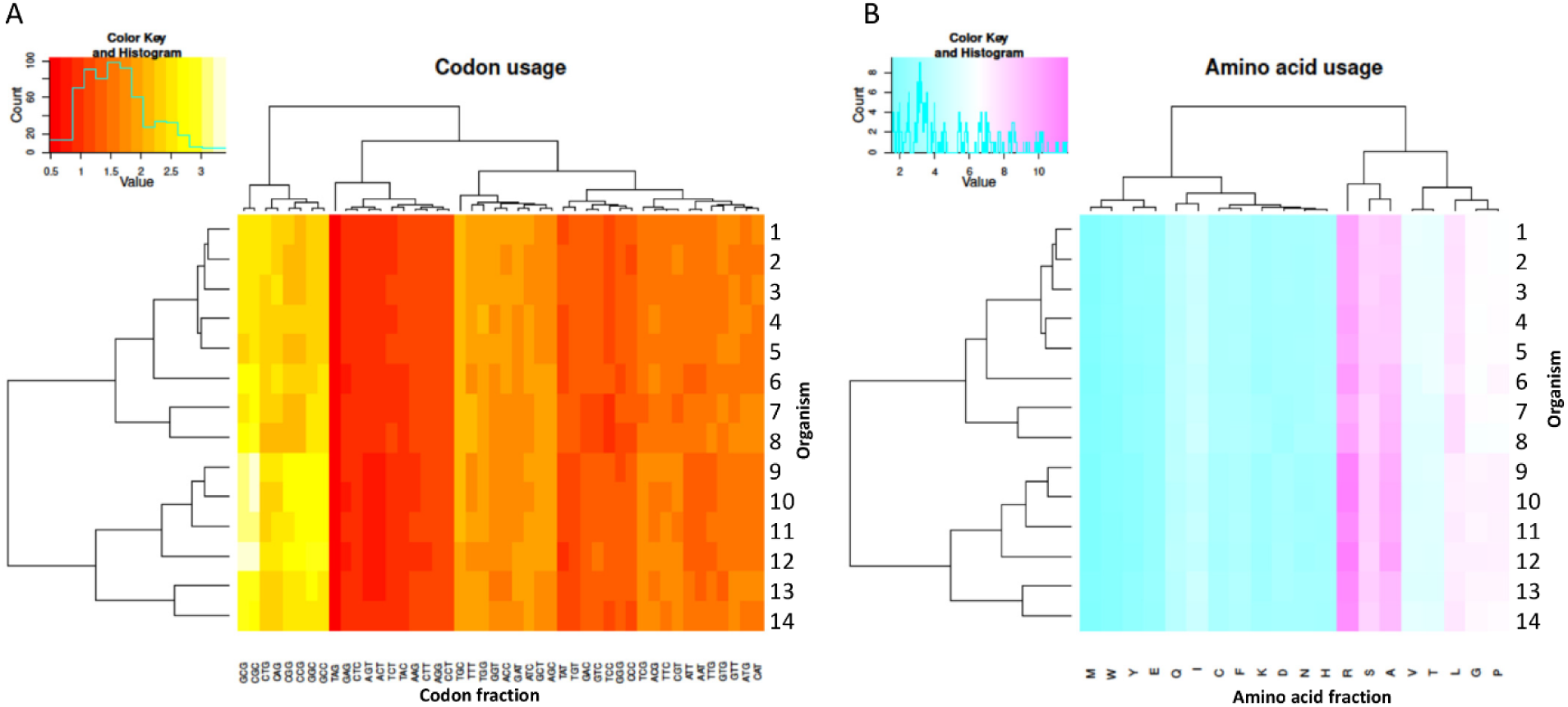
Amino acid and codon usage for all 14 genomes calculated based on the genes identified. The percentage of codon (**A**) and amino acid usage (**B**) were plotted in two heatmaps using R. Organisms were marked 1 to 14; 1. PL65, 2. A5410, 3. MS2, 4. Ech586, 5. EC1 (*D. zeae*), 6. *D. undicola* FVG10-MFV-A16, 7. *Dickeya aquatica* 174/2, 8. *D. lacustris* S29, 9. *D. solani* IPO 2222, 10. *D. dadantii* 3937, 11. *D. dianthicola* ME23, 12. *D. fangzhongdai* PA1, 13. *D. paradisiaca* Ech703 and 14. *D. chrysanthemi* Ech1591.

The intensity of color gradually changed from pink to blue when the amino acid frequency increased. The clades were distinct in generated phylogenetic patterns. The *D. zeae* species complex, *D. undicola, D. aquatica,* and *D. lacustris* were grouped together in the same clade while the remaining species formed a distinct lineage. Interestingly, *D. chrysanthemi* was phylogenetically distinct from *D. zeae* complex. The genomes of *D. paradisiaca* and *D. chrysanthemi* were together within the same group. Furthermore, in disagreement with the previous phylogenetic analysis, the amino acid and codon dendrograms displayed different relationships within the *Dickeya* species.

### 3.4. The pan and core genome analysis in *Dickeya genus*

To complement our previous analysis and obtain a better understanding of general genomic content similarities, a pan-core genome analysis was carried out among the fourteen *Dickeya* genomes. A pan-core genome plot is the corresponding calculated output and is presented in **Figure 5A**. A final core genome of 2,306 gene families, and a pan-genome of 9,450 gene families were obtained. The addition of genomes in the analysis caused a decrease in the core genome size, thus indicating a genome heterogeneity among the 14 *Dickeya* species. Considering the average gene number of ∼4,720 for the *Dickeya* genomes, 2,306 core genes represented approximately 50% of the total genome. As a result of this analysis, approximately half of the genomic constituents were conserved or orthologous across the 14 genomes analyzed in this study. Hence, the core size (2,306) pinpoints crucial genes potentially essential for *Dickeya* species survival. The high pan-genome size, which is more than three times the number of core size, seems to suggest significant genetic variation and evolutionary dynamics among the *Dickeya* species. The *D. zeae* complex strains showed the similar pattern of core- and pan-genomes size variation in **Figure 5A**, indicating the heterogeneity within this complex. To understand the genome-based relationships across the 14 *Dickeya* species, a phylogenetic tree was constructed using 2,306 conserved genes. The generated dendrogram formed a clear clade for *D. zeae* complex strains (**Figure 5B**).

**Figure 5.**
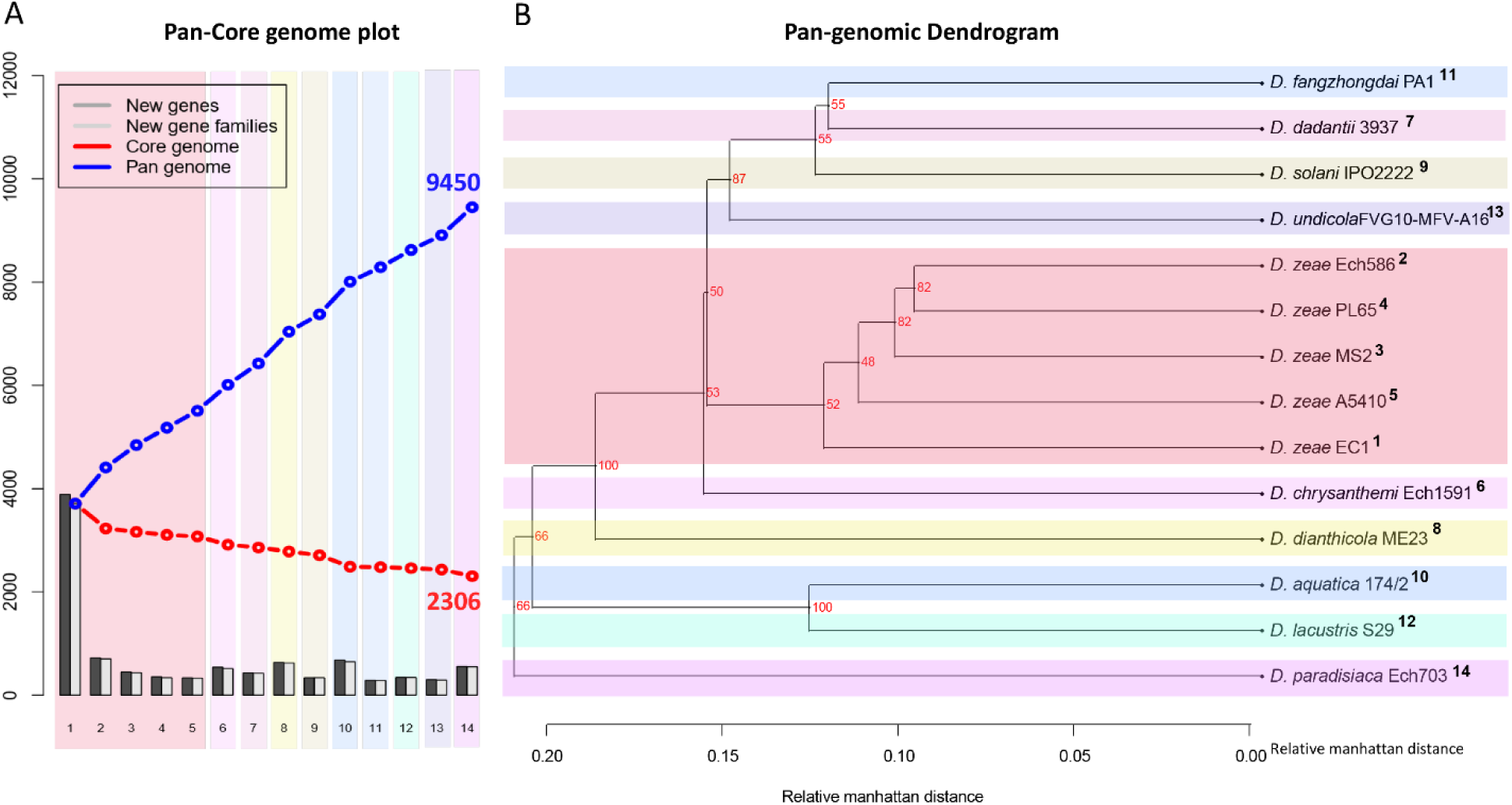
Pan- and core-genome analyses among the *Dickeya* species. Pan- and core-genome plot: The pan and core genome analyses were developed employing BLAST with a cutoff of 50% identity and 50% coverage of the longest gene (**A**). Pan-genome tree: the respective dendrogram illustrates the grouping between species based upon the shared gene families (core genome) defined in the pan and core genome analysis (**B**). Organisms are marked 1 to 14.

### 3.5. Comparison of main DNA features among *D. zeae* complex strains isolated from distinct hosts

To evaluate and compare the genomic properties of strains A5410 and PL65 sequenced in this study, BLAST atlases were created using these two genomes as references and compared with the other *D. zeae* complex strains EC1, Ech586 and MS2 (**Figure 6A-B**). The main DNA features namely: genome size, percent AT (red indicates high AT), GC skew (blue indicates most G’s prevalence), direct and inverted repeats (blue and red, respectively), position reference, stacking energy and intrinsic curvature were drawn in the atlas for each reference. Following these layers, all genome queries were displayed in the atlases as specific-colored lines, where only gene regions that matched with the reference organism were drawn. Several notorious divergences in the intrinsic genomic features of these two new strains were observed with respect to the other *D. zeae* strains. Strain A5410, for instance (**Figure 6A**), harbored 11 inverted and 20 direct repeats, and among them, 10 direct repeats were exclusive to this strain. Regarding the DNA properties, the strains A5410 (pineapple host) and PL65 (taro host) and Ech586 contained a genomic region that displayed low intrinsic curvature, low stacking energy and low position reference (pinpointed in dark red arrow). Moreover, three regions of low intrinsic curvature, low stacking energy but high position reference (pinpointed with dark blue arrow) were only found in the pineapple strain A5410. Another gene zone with the same DNA features was absent in strains MS2 and Ech586 (pinpointed with purple arrow). Interestedly, nineteen regions associated with high values of intrinsic curvature, stacking energy and position reference were merely observed in A5410.

**Figure 6.**
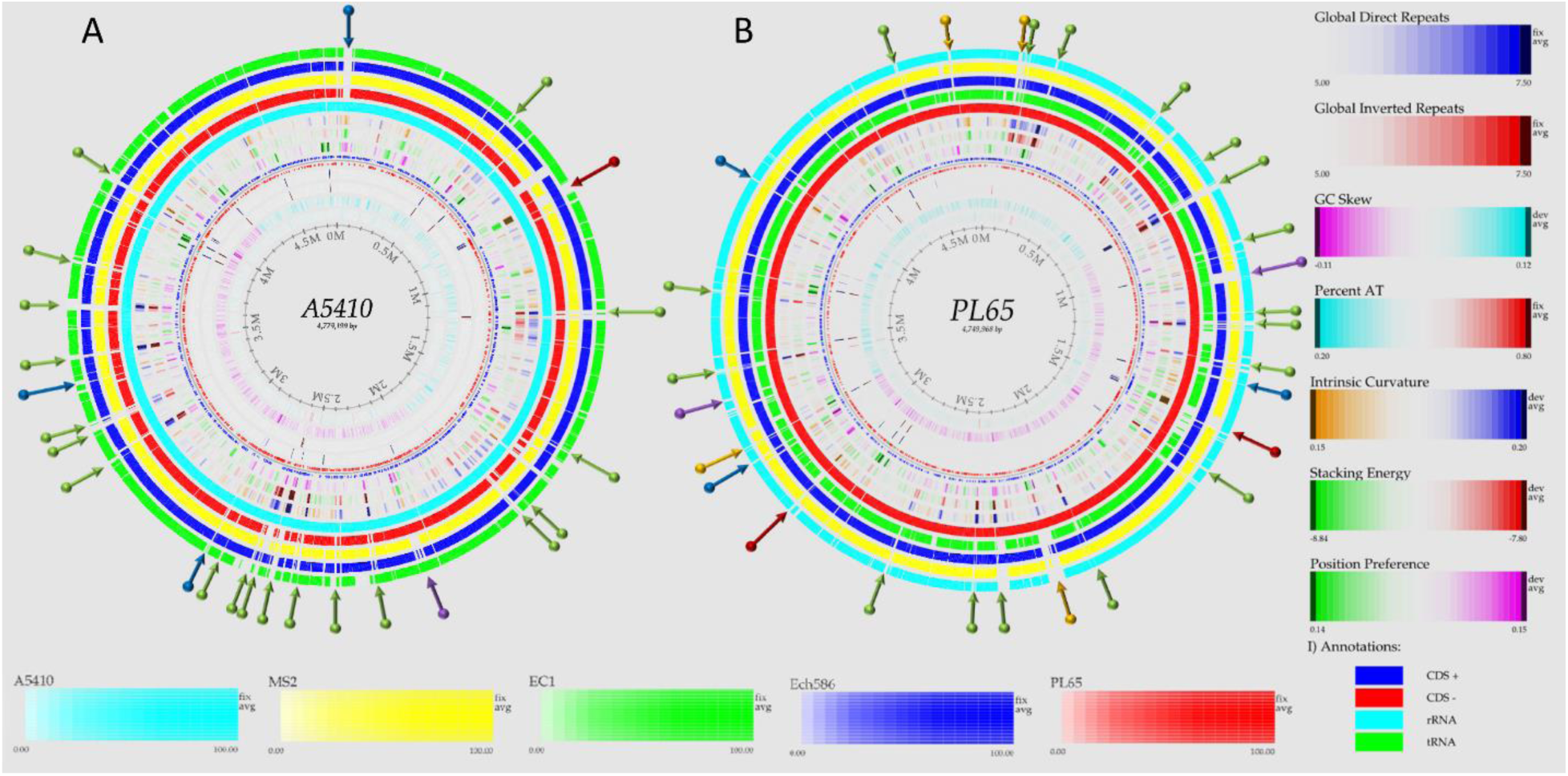
Genome BLAST atlases within the *D. zeae* complex species. The genomes of novel strains A5410 (**A**) and PL65 (**B**) were used as references to generate the circular graphics. Data of DNA, RNA and gene features of both references were obtained after annotating the genomes using the NCBI Prokaryotic Genome Annotation Pipeline (PGAP). From the inwards lane, the figures display the size of the genome (axis), percent AT (red = high AT), GC skew (blue = most G’s), inverted and direct repeats (color = repeat), position preference, stacking energy and intrinsic curvature. Following these layers, the external solid rings (indicated with unique colors) represent the genomes of other *D. zeae* strains mapped against the references. Olive arrows highlight those unique DNA regions associated with high intrinsic curvature, stacking energy and position reference found solely in the novel strains A5410 and PL65. Dark-red arrows pinpoint areas of the genome with low intrinsic curvature, stacking energy and position reference. Dark-blue arrows indicate low intrinsic curvature, low stacking energy but high position reference. Orange arrows show areas of the genome displaying high intrinsic curvature, high stacking energy but low position reference whereas purple arrows represent those genetic zones absent in some *D. zeae* isolates. BLAST genome atlases were created using the CMG-Biotools pipeline (Vesth et al. 2013).

On the other hand, the taro strain PL65 exhibited 12 and 22 inverted and direct global repeats, respectively (**Figure 6B**), of which 6 direct repeats were not found in the other isolates. This strain was the only one which presented three DNA regions with low intrinsic curvature, low stacking energy but high position reference features (pinpointed with arrow blue). Other two regions with similar features (pinpointed with purple arrow) were also observed; one of these regions was not observed in the genomes of strains A5410 and EC1 while the other was not present in MS2 and Ech586. Additionally, the strain PL65 exhibited four characteristic regions that displayed high intrinsic curvature, high stacking energy but low position reference values (pinpointed with orange arrows). These regions were not visualized in the other strains. Lastly, a total of seventeen gene regions with the same features mentioned previously but with a high position reference were shown by the genome of PL65 but not in the other organisms. Altogether, the DNA properties analyzed in these two new strains clearly demonstrated high divergence in the genome properties among the *D. zeae* strains.

### 3.6. General genomic features of two novel strains (PL65 and A5410)

The depth (X) of the assemblies were 558 and 701 for A5410 and PL65 strains, respectively. The complete genomes of novel strains, A5410 and PL65, consisted of a single circular chromosome of 4,501,560 and 4,501,560 base pairs, with a GC content of 54.6% and 54.6%, respectively.

The genome PL65 contains 4,182 protein-coding DNA sequences (CDS), 75 tRNA-coding genes, 22 ribosomal RNA-coding (5S-16S-23S) genes, nine non-coding RNA genes, and 87 pseudogenes. The genome A5410 contains 4,305 protein-coding DNA sequences (CDS), 75 tRNA-coding genes, 22 ribosomal RNA-coding (5S-16S-23S) genes, 8 non-coding RNA genes and 90 pseudogenes. Detailed information of five genomes is provided in **Table 2**. The length of *D. zeae* genomes were between 4.2 to 4.3 Mb. The GC content percent was almost similar (∼53%) for all five genomes. Regarding total conserved protein-coding sequence genes (CDS), the A5410 *D. zeae* genome displayed the highest CDS with 4,305 genes and highest pseudogenes with 90 genes (**Table 2**).

**Table 2.** General genome characteristics described for the five complete genomes of *Dickeya zeae* complex.

| Genome features | EC1 | Ech586 | MS2 | A5410 | PL65 |
| --- | --- | --- | --- | --- | --- |
| Length (bp) | 4532364 | 4818394 | 4740052 | 4779199 | 4749968 |
| % GC | 53.4 | 53.6 | 53.4 | 53.5 | 53.6 |
| CDS (coding) | 4012 | 4269 | 4221 | 4305 | 4182 |
| rRNAs (5S, 16S, 23S) | 8, 7, 7 | 8, 7, 7 | 8, 7, 7 | 8, 7, 7 | 8, 7, 7 |
| tRNAs | 88 | 76 | 75 | 75 | 75 |
| ncRNAs | 12 | 6 | 6 | 8 | 9 |
| Pseudogenes | 65 | 64 | 57 | 90 | 87 |
| Hypothetical proteins* | 760 | 565 | 924 | 879 | 873 |
| Virulence factor* | 71 | 72 | 71 | 69 | 69 |
| Transporter* | 176 | 180 | 186 | 183 | 183 |
| Antibiotic resistance* | 37 | 37 | 42 | 42 | 42 |
| Drug target [TTD]* | 27 | 27 | 26 | 27 | 27 |

GC, Guanine-Cytosine; CDS, Coding Sequence Regions; EC number, Enzyme Commission Number; TTD, Therapeutic Target Database.
\*Data retrieved from the bioinformatics PATRIC webserver.

OrthoMCL pipeline was used to develop a robust comparative genomics analysis. The results showed that the number of core genes among the five genomes were 3,162 genes. The number of specific genes in A5410, PL65, Ech586, EC1, and MS2 were 137, 102, 123, 143, and 158, respectively (**Figure 7**).

**Figure 7.**
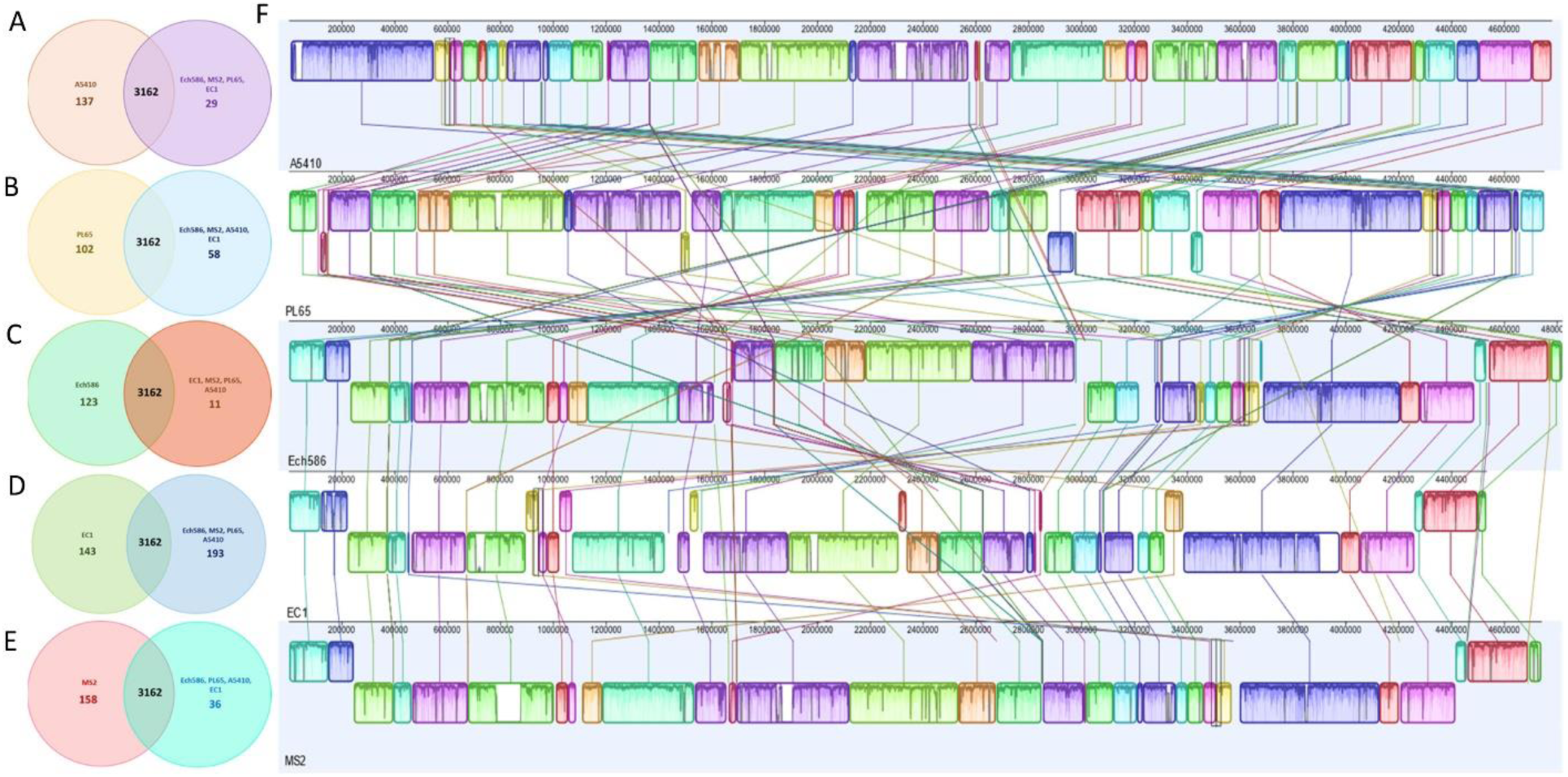
Venn diagrams (**A**), (**B**), (**C**), (**D**) and (**E**) for the deduced proteins of A5410, PL65, Ech586, EC1 and MS2, respectively. Values were calculated by OrthoMCL clustering analyses using the following parameters: P-value Cut-off = 1 × 10−5; Identity Cut-off = 90%; Percent Match Cut-off = 80. The overlapping sections indicate shared numbers of deduced proteins. In (**F**), pairwise alignments of five linearized *D. zeae* genomes were generated using Mauve.

The genomes of two novel strains were analyzed extensively. The genome A5410 harbored some essential unique genes, such as transporter proteins, endonuclease protein, phage tail protein, aspartate/glutamate racemase family protein *Ars*D gene involved in arsenic resistance (Firrincieli et al., 2019), the carboxymuconolactone decarboxylase protein involved in protocatechuate catabolism (Cheng et al., 2017). The nitrogen fixation gene cluster was only present in A5410 strain isolated from pineapple (**Figure 8A**). The genome PL65 harbored some essential unique genes, such as glycosyltransferase protein, transporter proteins, endonuclease protein, and phage tail protein. The pilus assembly protein cluster was only present in PL65 genome isolated from taro (**Figure 8B**).

**Figure 8.**
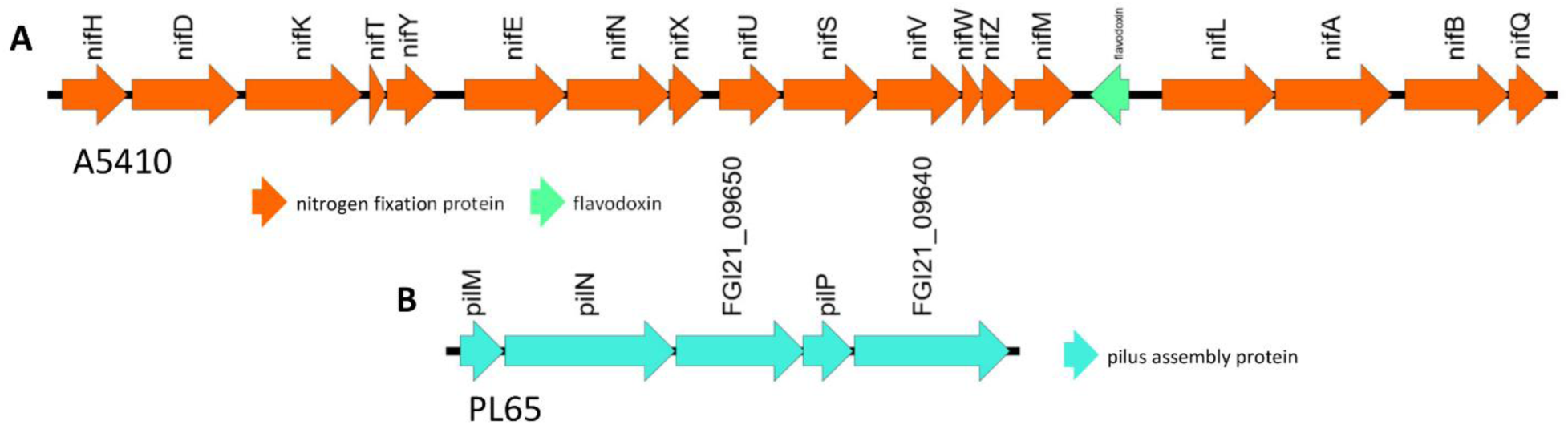
The unique gene clusters. The Figure (**A**) shows the nitrogen fixation cluster from the genome A5410. The Figure (**B**) shows the pilus assembly protein cluster from PL65.

Significantly, pilus cluster was predicted within a genomic island (GI) (**Figure 9**). Genomic islands (GIs) or horizontal acquired islands (HAIs) are incorporated into the bacterial genome during the conjugation process, and besides, harbor genes are required for integration and excision into the chromosome (Johnson and Grossman, 2015; Zakharova and Viktorov, 2015). The genomes of PL65 and A5410 were screened for horizontally acquired DNA using IslandPath-DIMOB, SIGI-HMM, IslandPick, and Islander methods integrated in the IslandViewer server (Bertelli et al., 2017) (**Figure 9**).

**Figure 9.**
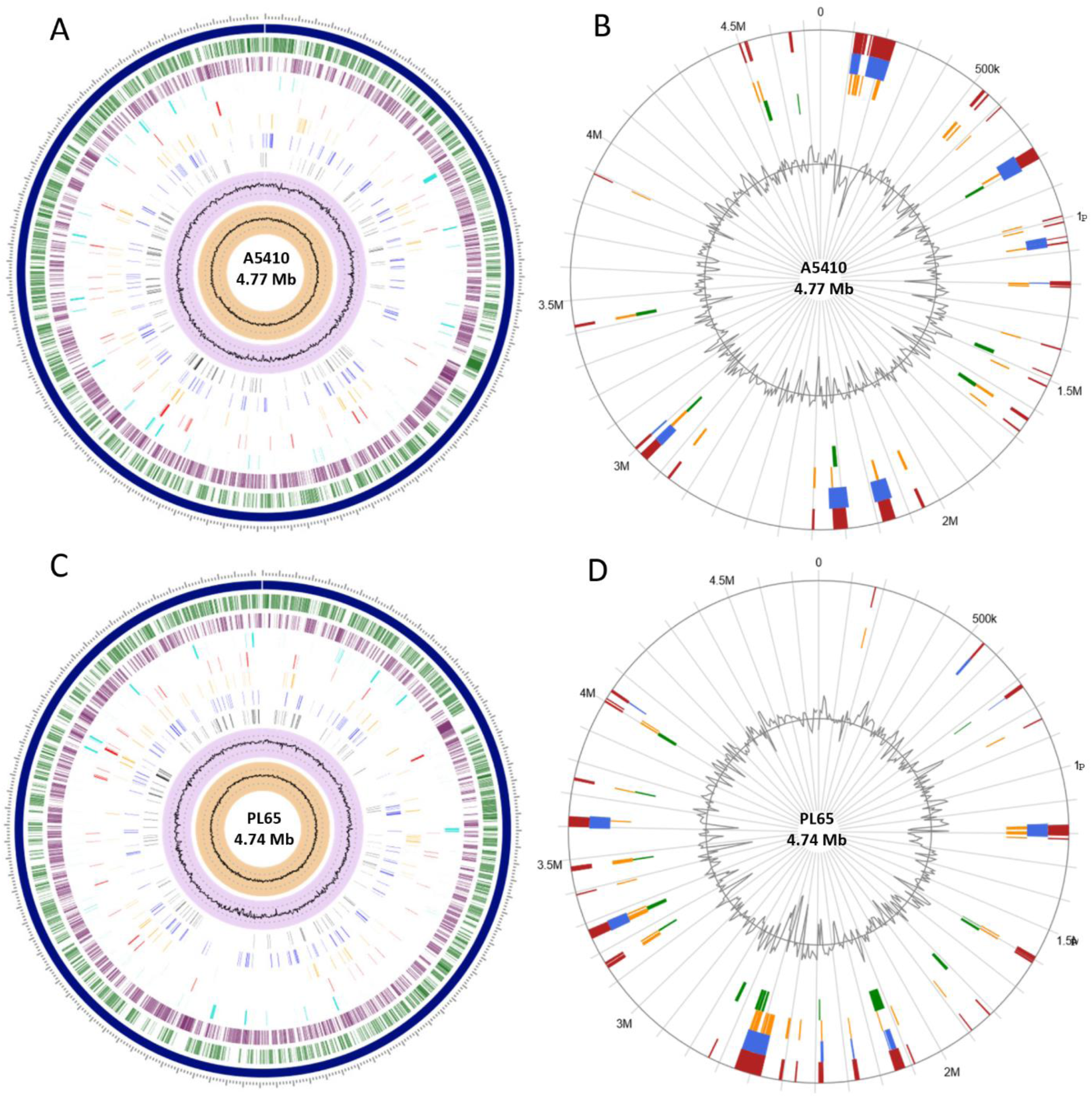
Circular view of the genomes of A5410 (**A**) and PL65 (**C**) strains generated using PATRIC showing the physical map of significant features. From outside in: Position label Mb (shown in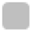); order of contigs (shown in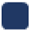); distribution of coding sequences in forward strands (shown in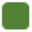); distribution of coding sequences in reverse strands (shown in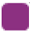); distribution of noncoding elements along the chromosome (shown in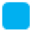); distribution of genes involved in antibiotic resistance (shown in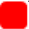); distribution of other virulence factors (shown in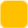); distribution of genes encoding transporter proteins (shown in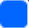); distribution of genes encoding drug targets (shown in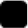); distribution of GC content (shown in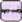); and distribution of GC skew (shown in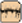). Circular visualization of predicted Genomic Islands (GIs) on A5410 (**B**) and PL65 (**D**) strains. The analysis was conducted in the IslandViewer 4. The interactive visualization of the distinct islands across the genomes are shown with blocks colored according to the predictor tool as described: IslandPick (shown in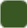) based on genome comparison, IslandPath-DIMOB (shown in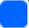) based on associated GIs features such as tRNAs, transposon elements, integrases and sequence bias, SIGI-HMM (shown in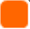), based on the codon usage bias with a Hiddden Markov model criterion and the integrated results of the four tools (shown in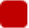).

In the genome A5410, 56 presumed genome islands (GIs) ranged 2.6 to 83.1 kb were detected. The largest GIs consisted of 83,100 bp with 92 predicted gene coding regions, while the shortest GIs consisted of only five predicted gene coding regions. A total of 707 genes were predicted into the GIs. Only 68 Open Reading Frames (ORFs) were unique genes for A5410 among analyzed genomes. For strain PL65, 47 presumed GIs that ranged 3.1 to 70.3 kb were detected, of which the largest consisted of 70,338 bp and predicted to encode 69 genes. A total of 675 genes were predicted into the GIs. Only 55 ORFs were unique genes for PL65 among analyzed genomes.

Genes encoding tRNAs, transposases and integrases, pectate lyase, endoglucanase, and phage tail protein and genes related to the citrate synthase system, the type VI secretion system (VgrG), the type IV secretion system (Rhs), toxin-antitoxin systems, and antibiotic biosynthesis were identified in the GIs of both strains. Furthermore, genes related to flagella and chemotaxis, type IV pilus biogenesis system, CRISPRcas1, CRISPRcas2, type III CRISPRCsm1-6 were identified in the GIs of strain PL65.

### 3.7. Genome comparison of gene clusters associated with pathogenesis among the *Dickeya zeae* complex

The soft rot bacteria within the genus *Dickeya*, macerate the plant tissue, and acquire nutrients from the dead cells (Davidsson et al., 2013). Several pathogenicity determinant genes are involved in this process. Most pathogenicity determinants genes including the plant cell wall degrading enzymes (PCWDE), type secretion systems (Type I-VI), synthesis of polysaccharides (enterobacterial common antigen, capsular polysaccharide, lipopolysaccharides, exopolysaccharides, and O-antigen), bacterial attachment operons (type IV pili), flagella and chemotaxis, quorum-sensing systems and zeamine synthesis have been described for EC1 strain (Zhou et al., 2015). We analyzed and compared the similarities, differences, or absence of virulent determinant genes among the *D. zeae* complex. Studies were carried out regarding the genome of EC1 strain.

#### 3.7.1. Plant cell wall-degrading extracellular enzymes and proteases

The PCWDE, including pectinases, polygalacturonases, cellulases, and proteases, are essential virulence determinants, which degrade the structural components of the plant cell wall, playing a significant role during bacterial pathogenesis and disease development (Toth et al., 2006; Charkowski et al., 2012; Nykyri et al., 2012; Davidsson et al., 2013). The PCWDE are the main responsible factors in creating soft rot symptoms. Because of that, we focused on this factor. All *Dickeya* species were analyzed and compared for PCWDE.

Many of these pectinases—pectate lyase (*Pel*), pectin lyase (*Pnl*), pectin methyl esterase (*Pme*) and polygalacturonase (*Peh*)—are scattered throughout the genomes rather than arranged in clusters (Glasner et al., 2008; NyKyri et al., 2012). The pectinases encoded by independent genes seem to be derived from successive rounds of gene duplicated (Barras et al., 1987; McMillan et al., 1994). *Pel*ABCDEILNZ genes encoding endo-pectate lyases and *Pel*W and *Pel*X genes encoding pectate disaccharide-lyases were highly conserved in *D. zeae* genomes and other *Dickeya* species, except that *Pel*D was absent in *D. dianthicola* ME23, *Pel*E was absent in *D. lacustris* S29, *D. paradisiaca* Ech703, and *D. aquatica* 174/2, and *Pel*I was not found in *D. paradisiaca* Ech703 (**Table S1 and** **Figure 2B**). Similarly, *Pem*AB genes encoding pectin methylesterases were conserved in *Dickeya* species. However, *Pem*B is absent in *D. zeae* genomes, *D. lacustris* S29, *D. paradisiaca* Ech703, and *D. aquatica* 174/2 (**Table S1 and** **Figure 2B**).

The *Peh*KNVWX genes encoding polygalacturonase were also conserved in *Dickeya* species. Surprisingly, the *Peh*VWX genes were identified with 99% similarity (**Table S1 and** **Figure 2B**). The *PehX* has two copies in *D. dianthicola* ME23 (*PehV* and *PehX*) and three copies in *D. dadantii* 3937 and *D. solani* IPO2222 (*PehV*, *PehW*, and *PehX*). Moreover, the *pnl*GH genes encoding pectin lyase were present in *D. dadantii* 3937 and *D. dianthicola* ME23. However, *D. zeae* Ech586, A5410, and PL65 indicated the loss of these genes (**Table S1 and** **Figure 2B**). All species harbored the pectin acetylesterases (*PaeX* and *PaeY*) (**Table S1 and** **Figure 2B**). The oligogalacturonate lyase (*Ogl*) was present in all analyzed *Dickeya* species (**Table S1 and** **Figure 2B**). The *Gan*ABCEFGKLR gene cluster, responsible for removing galactan chains in pectin ramified regions, were present in all *Dickeya* species (Hugouvieux-Cotte-Pattat et al. 2014). However, the

MS2 strain of *D. zeae* and *D. paradisiaca* Ech703 lost the *Gan*ABCEFG genes (**Table S1 and** **Figure 2B**). The rhamnogalacturonate lyase *Rhi*E gene was present in all species except *D. paradisiaca* Ech703 (**Table S1 and** **Figure 2B**). The ferulate esterase *Fae*D and *Fae*T genes were present in all *Dickeya* species except *Fae*T gene, which was lost in *D. zeae* EC1 and A5410 (**Table S1 and** **Figure 2B**). The regulator *Kdg* was present among the *Dickeya* species. However, the *Kdg*T transporter gene was absent in *D. lacustris* S29 and *D. aquatica* 174/2; the *Kdg*N transporter gene was absent in *D. paradisiaca* Ech703 (**Table S1 and** **Figure 2B**). The *Exu*RT, *Kdu*DI, *Uxa*ABC genes, and *Tog*ABMNT transporter were highly conserved and present in all tested genomes (**Table S1 and** **Figure 2B**).

#### 3.7.2. Cellulases and xylanases enzymes

The endoglucanase genes (*cel*Y and *cel*Z), beta-glucosidase encoding genes (*bglA*, *bgxA*, *bglB*, *nagZ*, *bglC*, *bglD,* and *celH*) and an alpha-glucosidase encoding gene (*lfaA*) are involved in the degradation of cellulose to glucose (Zhou et al., 2015). Although these genes were conserved in the *Dickeya* genus, the *bglC* gene was absent in some *D. zeae* genomes (MS2, Ech586, and A5410), *D. lacustris* S29, and *D. aquatica* 174/2 (**Table S1 and** **Figure 2B**). The *bgl*D gene was also absent from the *D. zeae* genomes MS2 and Ech586, *D. lacustris* S29, and *D. aquatica* 174/2 (**Table S1 and** **Figure 2B**). The *lfaA* gene was not present in *D. lacustris* S29. The xylanases (*XynA*) gene is involved in the degradation of xylan and xyloglucan, mainly present in plant cell walls (Pena et al., 2016). *Dickeya zeae* genomes (A5410, PL65, EC1, Ech586, and MS2), *D. dadantii* 3937, *D. paradisiaca* Ech703, *D. fangzhongdai* PA1, and *D. solani* IPO 2222 contained *Xyn*A (**Table S1 and** **Figure 2B**). Interestingly, the xylose degradation enzymes (*Xyl*ABFGHR) were present in all *Dickeya* species except *D. lacustris* S29 and *D. aquatica* 174/2.

#### 3.7.3. Type secretion systems (TSS)

The type I secretion system (T1SS) constitutes *prt*D, E, and F, and responsible for secreting the proteases. The *prt* cluster encodes four metalloproteases (*prt*ABCG) and three protease secretion associated proteins (*prt*DEF) (Zhou et al., 2015). The metalloprotease *prt*W plays an important role to degrade the plant cell wall proteins (Charkowski et al., 2012; Zhou et al., 2015). On the other hand, the Type II secretion system (T2SS) is responsible for translocating extracellular proteins across the outer membrane (Jha et al., 2005, Zhou et al., 2015). The *out* cluster eencodes two outer membrane proteins (*out*SD), five inner membrane proteins (*out*BEFLM), one trans periplasmic protein (*out*C), and prepilin peptidases (*out*GHIJLK) (Zhou et al., 2015). T1SS (within the range of 66-100% nucleotide identity) and T2SS (within range of 82-100% nucleotide identity) were present and highly conserved in the *D. zeae* genomes [Locus tag of T2SS; EC1: W909_RS13390-RS13455, Ech586: DD586_RS13945-RS14010, MS: C1030_RS14365- RS14430, A5410: FGI04_04420-04355, and PL65: FGI21_21175-21110. Locus tag of T1SS; EC1: W909_09760-09795, Ech586: DD586_2059-2052, MS: C1030_10785-10820, A5410: FGI04_08210-08175, and PL65: FGI21_03340-03305] (**Figure S3 A-B**).

The Type III secretion system (T3SS) is integrated by the *hrp* (hypersensitive response and pathogenicity) and *hrc* (hypersensitive response conserved) genes clusters (Toth et al., 2006, Zhou et al., 2015). According to our analyses, T3SS was present and highly conserved within a range of 85-100% nucleotide identity in the genomes of *D. zeae* complex (EC1, MS2, Ech586, and A5410), except PL65 [Locus tag of T3SS; EC1: W909_RS10510-RS10380, Ech586: DD586_RS09320- RS09470, MS: C1030_RS11710-RS11560, A5410: FGI04_007220-07360, and PL65:

FGI21_02600-02615] (**Figure 10**). Most genes of T3SS were absent in PL65 except for the *Hrp*E (FGI21_02600) and *Hrp*U (FGI21_02590) genes. While the size of *Hrp*U was 1,080 bp, the *Hrp*U gene of PL65 is only 158 bp, with an average 94% identity. The function of *Hrp*E was discovered as an elicitor of plant defense responses (Gottig et al., 2018). The *Hrp*E gene was described as a primary structural component of T3SS (Gottig et al., 2018). The *D. zeae* (EC1, MS2, Ech586, and A5410) genomes harbored a large gene cluster with three transcriptional units identified, spanning a genomic region of approximately 25 kb. The genetic region of T3SS was between the *hrp*N and *hrc*U genes within the genomes of *D. zeae* complex. The *plc*A gene, which encodes an extracellular phospholipase, was present in all *D. zeae* genomes. The *hrp* and *hrc* gene clusters and the near-upstream region of the *plc*A gene were highly conserved. While there were no additional genes in the EC1 genome, MS2 and Ech586 genomes harbored four extra ORFs encoding three hypothetical proteins and a membrane-bound lytic murein transglycosylase *Mlt*B. Additionally, A5410, and PL65 genomes harbored two ORFs encoding one hypothetical protein and *mlt*B gene (**Figure 10**).

**Figure 10.**
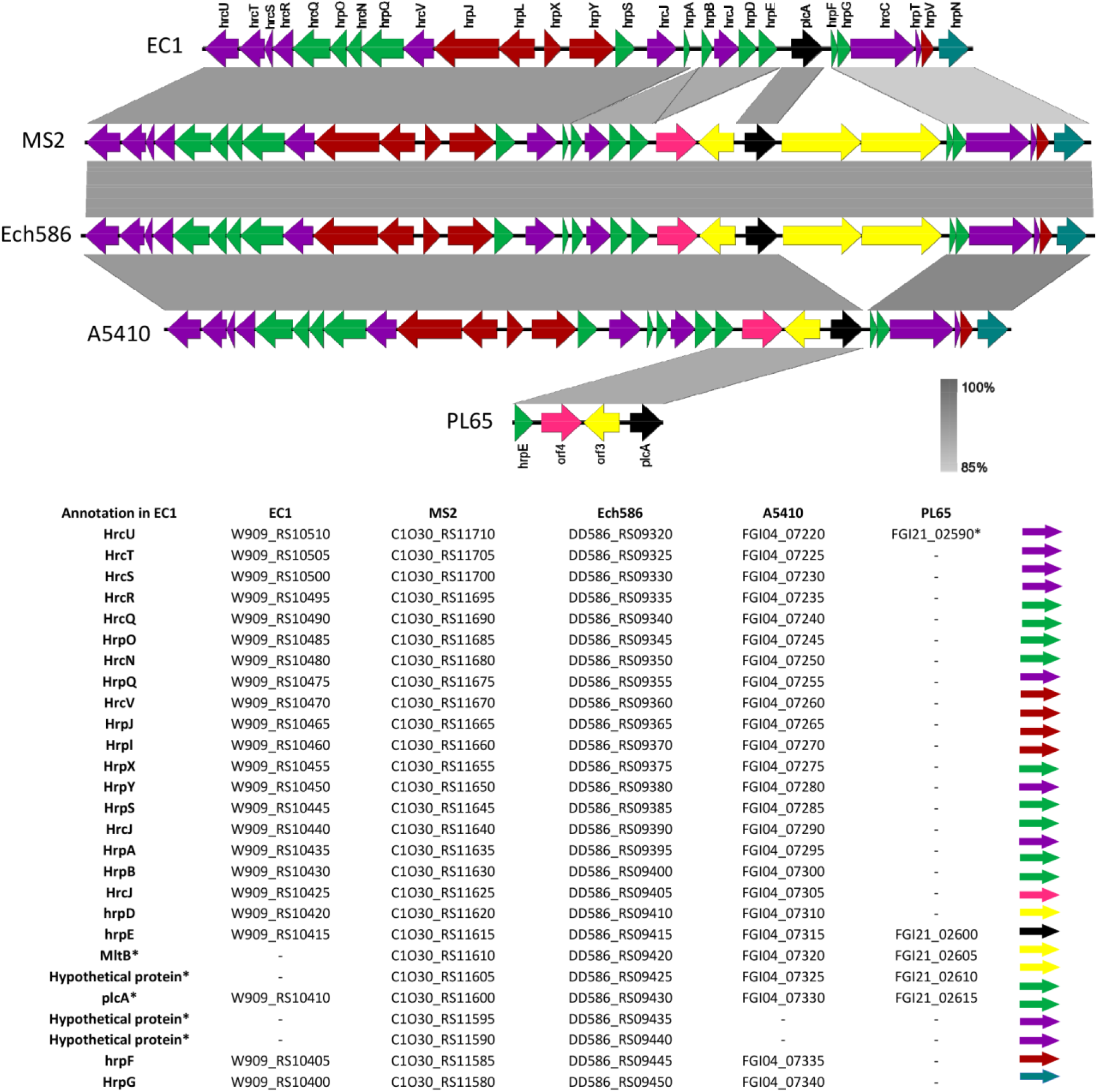
Comparison of the genetic organization of type III secretion system among five *Dickeya zeae* strains. The arrow position represented forward/reverse gene orientation. Arrow color signified specific gene composition within the T3SS. A pairwise alignment between the linear sequences was rendered based upon BLAST algorithm with cut-off values from 85% to 100%. Regions with higher nucleotide identity were displayed with a shaded grey.

Type IV secretion system (T4SS) constitutes virB1-11 genes and functions in conjugation, pathogenicity, and DNA release/uptake (Bhatty et al., 2013; Trokter et al., 2014; Darbari et al., 2015). The T4SS gene cluster is highly conserved within the range of 91-100% nucleotide identity among EC1, Ech586, and A5410, but not found in PL65 and MS2; both strains lacked most of T4SS genes and only had virB1 and virB2 (**Figure 11**).

**Figure 11.**
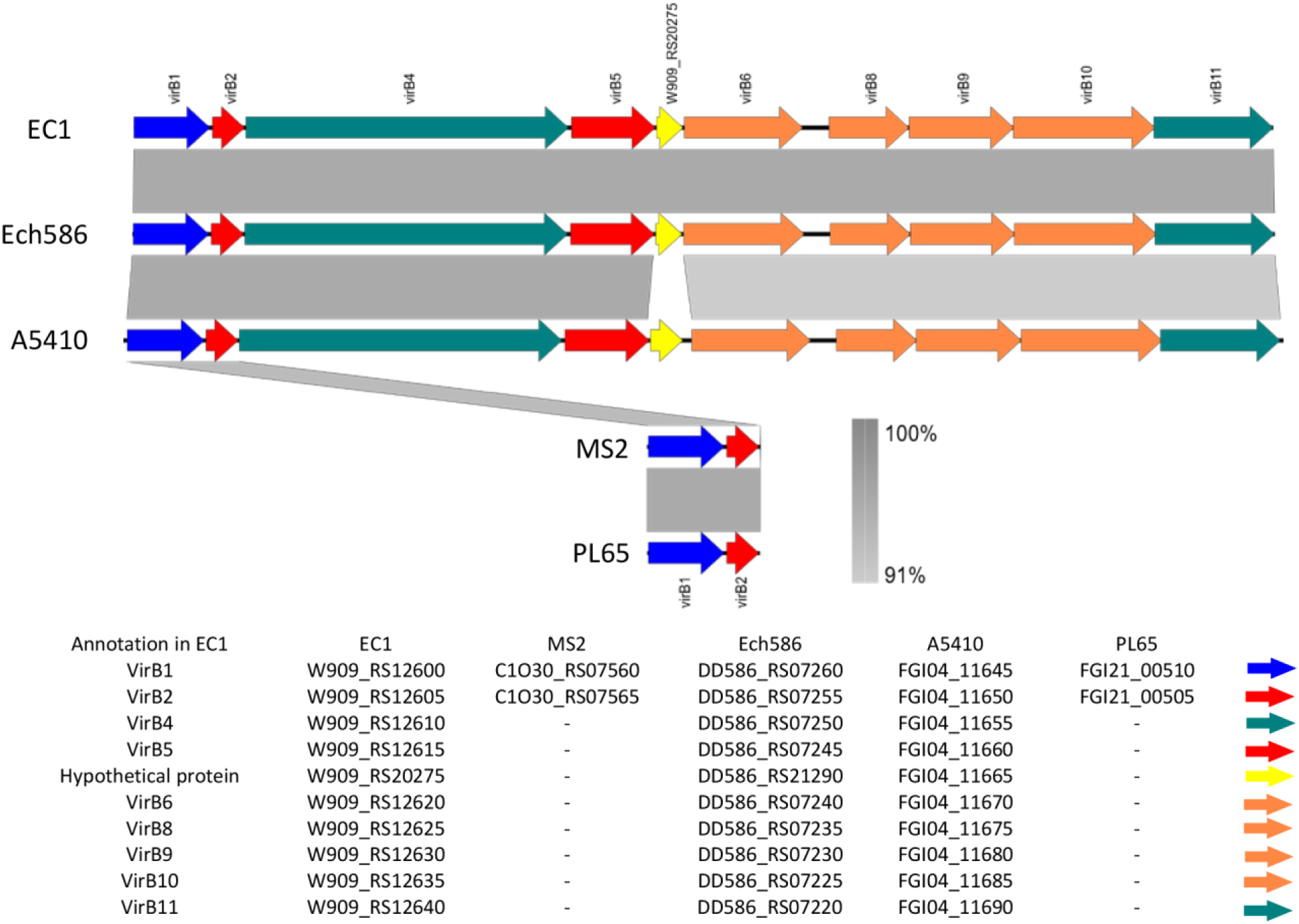
Comparison of the genetic organization of type IV secretion system among five *Dickeya zeae* strains. The arrow position represented forward/reverse gene orientation. Arrow color signified specific gene composition within the T4SS. A pairwise alignment between the linear sequences was rendered based upon BLAST algorithm with cut-off values from 91% to 100%. Regions with higher nucleotide identity were displayed with a shaded grey.

The T5SS encompasses either one or two proteins, the latter constituting two-partner secretion systems called *Hec*/*Tps*/*Cdi* (contact-dependent inhibition) (Charkowski et al., 2012; Pédron et al., 2014). In the *D. zeae* genomes, just one *Hec*/*Tps*/*Cdi* system waw present and harbor one *hec*B and one *hec*A genes that have been shown to act in contact-dependent growth inhibition (CDI) by delivering the C-terminal toxin domain of HecA (TspA/CdiA) to target cells (Pedron et al., 2014). The genes hecA2 and hecB (T5SS) were located near to the T3SS in all *D. zeae* complex. The hemagglutinin-coding loci *hecB* and *hecA* were present within the range of 68-100% nucleotide identity among the *D. zeae* complex strains [Locus tag of T5SS; EC1: W909_RS10375, W909_RS19830, Ech586: DD586_RS09475-RS22000, MS: C1O30_RS11550-RS11555, A5410: FGI04_07365-07405, and PL65: FGI21_02620-02625] (**Figure S4 A**). All *D. zeae* genomes harbored a *hec*B homolog. The *hec* system was universal among these *D. zeae* genomes. However, we found that the gene sequences encoding *hec*A2 in some *D. zeae* genomes were truncated. For instance, the gene sequences encoding *hec*A2 in Ech586 and A5410 were incomplete, and the entire system was annotated as pseudogenes. This finding indicated that this cluster might not be functional in these genomes. Besides, we observed inserted genes in A5410 and Ech586 genomes. One hypothetical protein and one type I toxin-antitoxin (TA) system protein (*Sym*E) were located between the first and second ORF for *Hec*A protein in Ech586 genome. Additionally, A5410 harbored six extra genes: three hypothetical genes, two *Sym*E, and one membrane channel gene— between incomplete *Hec*A genes. *Hec*A protein has two ORFs, and between those ORFs, three hypothetical proteins were located, two types I toxin-antitoxin (TA) system protein (*Sym*E) and one IS3 (Insertion sequences) transposase. The genome of EC1 harbored *hec*A gene; however, *hec*A exhibited a 50% query cover and 94% identity with PL65 and MS2 genomes.

The type VI secretion system (T6SS) targets other bacteria and thus plays an essential role as a polymicrobial injectosome that resembles as a bacteriophage tail (Bernard et al., 2011). The T6SS assists in multiple biological processes, for an instant, interaction with host eukaryotic cells, pathogenicity, antibacterial activity, symbiosis, metal ion acquisition, and biofilm formation (Bernard et al., 2011; Cianfanelli et al., 2016; Gallique et al., 2017). Regarding to the *D. zeae* complex genomes, the T6SS cluster was confirmed by 17 genes. The T6SS consists of the *hcp*, *vgr*G (virulence-associated protein G), *imp*BCF, and *vas*ABCDEFGHIJKL genes. The T6SS genes were highly conserved within the range of 68-100% in EC1, MS2, Ech586, A5410, and PL65 [Locus tag of T5SS; EC1: W909_RS06255-RS06425, Ech586: DD586_RS06380-RS06525, MS: C1O30_RS06775-RS06960, A5410: FGI04_12385-12215, and PL65: FGI21_07465-07265].

Additionally, an average of 20 genes were inserted into the T6SS cluster in all five genomes (**Figure S4B**). We found substantial variations in the extra set of genes inserted between *vgr*G and *imp*B (**Figure S4B**). The inserted cluster was annotated as ankyrin genes, hypothetical proteins genes, the repeat-containing protein *Rhs*As (rearrangement hot spot), amidohydrolase gene, *Sym*E genes, and *Par*DE genes.

#### 3.7.4. Flagellar and chemotaxis genes

The flagellar biosynthesis and chemotaxis clusters constitute 20 *Fli* genes (*Fli*ZACD-T), 14 Flg genes (*Flg*A-N) and 5 *Flh* genes (*Flh*A-E), involved in the flagellar synthesis, four flagellar rotation genes (*Mot*AB, MCP1, and MCP3) and six chemotaxis-associated genes (*Che*ABZYRW) (Zhou et al., 2015). The flagellar biosynthesis and chemotaxis genes were present and highly conserved in *D. zeae* genomes (**Table S2**). Interestingly, the EC1 genome harbored two sets of *Fli*C genes. There are 12 additional genes inserted between *Fli*A and *Fli*C in EC1, Ech586, and MS2 genomes. These inserted genes consisted two methyltransferases (*Rfb*C and *Fkb*M), an aminotransferase (*Sps*C), and nine fatty acid biosynthesis genes (*Ald*H, *Lux*E, *fad*D, *Tkt*AB two set of *Fab*G, *Acp*P, and *Maa*). Moreover, A5410 and PL65 also contained six inserted genes (*Rfb*C, two sets of *Fkb*M, *Car*A, and *Moc*A), which were highly conserved for A5410 and PL65 genomes (**Table S2**).

#### 3.7.5. Twitching motility genes

The type IV pilus biogenesis encoding system consist of *pil*F (type IV pilus biogenesis and stability protein), *Pil*T (Twitching motility protein) and *Pil*ABC, and *Pil*M-Qgenes (Maier and Wong, 2015; Duprey et al., 2019). The *pil*F and *Pil*T genes were located distant to the type IV pilus biogenesis cluster. The type IV pilus biogenesis system is present and highly conserved in the *D. zeae* genomes (**Table S2**).

#### 3.7.6. Polysaccharide genes

The ability to biosynthesize polysaccharides, which can be extracellularly secreted polysaccharide (EPS) or remain attached to the bacterial cell surface (lipopolysaccharide and capsular polysaccharides), is another essential factor for disease growth (Whitfield 2006). Bacteria display different types of polysaccharides, namely: lipopolysaccharide (LPS), which attach to the cell membrane, capsular polysaccharides that bind covalently to the cell surface, lipooligosaccharides (LOS) that lacks the O antigen, and the extracellular polysaccharides (EPS), which are secreted to the surrounding environment (Reeves et al., 1996). In our analysis, we observed that capsular polysaccharide (CPS) cluster was composed of 12 genes (*Cps*ABC-WcaB) which were highly conserved in Ech586, MS2, PL65, and A5410. However, the entire cluster was absent in EC1 (Table S2). The enterobacterial common antigen (ECA) composed of 10 genes (*Rff*M-*Wec*A) that were present in all genomes (**Table S2**). Interestingly, the genome MS2 harbors one inserted gene (hypothetical protein) between WzzE and WecA genes. The extracellular polysaccharides (EPS) cluster was composed of 22 genes (*Gnd*-*Wza*). All *D. zeae* genomes harbored the EPS cluster. However, EC1 and Ech586 had one extra glycosyltransferase protein located between *Gnd* and *Gal*F genes. The genome Ech586, A5410, PL65 harbored highly conserved lipopolysaccharide cluster (LPS), which was encoded by 11 genes (*Coa*D-*Rfa*D) (Table S2). In the case of the genome EC1, LPS cluster showed the high rearrangements (**Table S2**). The five genes of glycosyltransferase protein, including *Rfa*G and *Rfa*Q, were absent in EC1. However, EC1 genome harbored additional six glycosyltransferase genes and three hypothetical genes. Unlike the others, the LPS cluster of EC1 presented 14 genes. (**Table S2**)

#### 3.7.7. *D. zeae* complex contained different CRISPR-Cas Systems

Most bacteria harbor the CRISPR-Cas immunity systems to protect themselves from foreign genetic elements (Makarova et al., 2011). CRISPR-Cas systems contain two groups, class 1 (types I, III and IV), which includes interaction with multi-Cas protein complexes, and class 2 (types II, V and VI), which uses a single interaction effector protein (Hille et al., 2018). The CRISPR-Cas systems were identified in five *D. zeae* genomes using the CRISPRfinder online server (Grissa et al., 2007). *Dickeya zeae* strains contained three types of CRISPR-Cas systems (subtype I-F, subtype I-E, and type III-A) (**Table 3 and** **Figure 12**). The genome of EC1, Ech586, MS2, and PL65 harbored highly conserved subtype I-E CRISPR-Cas system composed of *cas*3, *cas*A, *cas*B, *cas*7e, *cas*5e, *cas*6e, c*as*1e, and *cas*2 (**Table 3**). The Ech586, MS2, A5410, and PL65 genomes presented the subtype I-F CRISPR-Cas system. The regular subtype I-F CRISPR-Cas system contains *cas*1f, *cas* 3f, *csy*1, *csy*2, *csy*3, and *cas*6f as found in MS2. However, Ech586, A5410 and PL65 confined a set of sequences downstream of c*as*6f operon. These genes were identified as coding sequences of the *Asn*C transcriptional regulator protein, *Yit*T integral membrane protein, aspartate/tyrosine/aromatic aminotransferase protein, amino acid permease and other hypothetical proteins.

**Figure 12.**
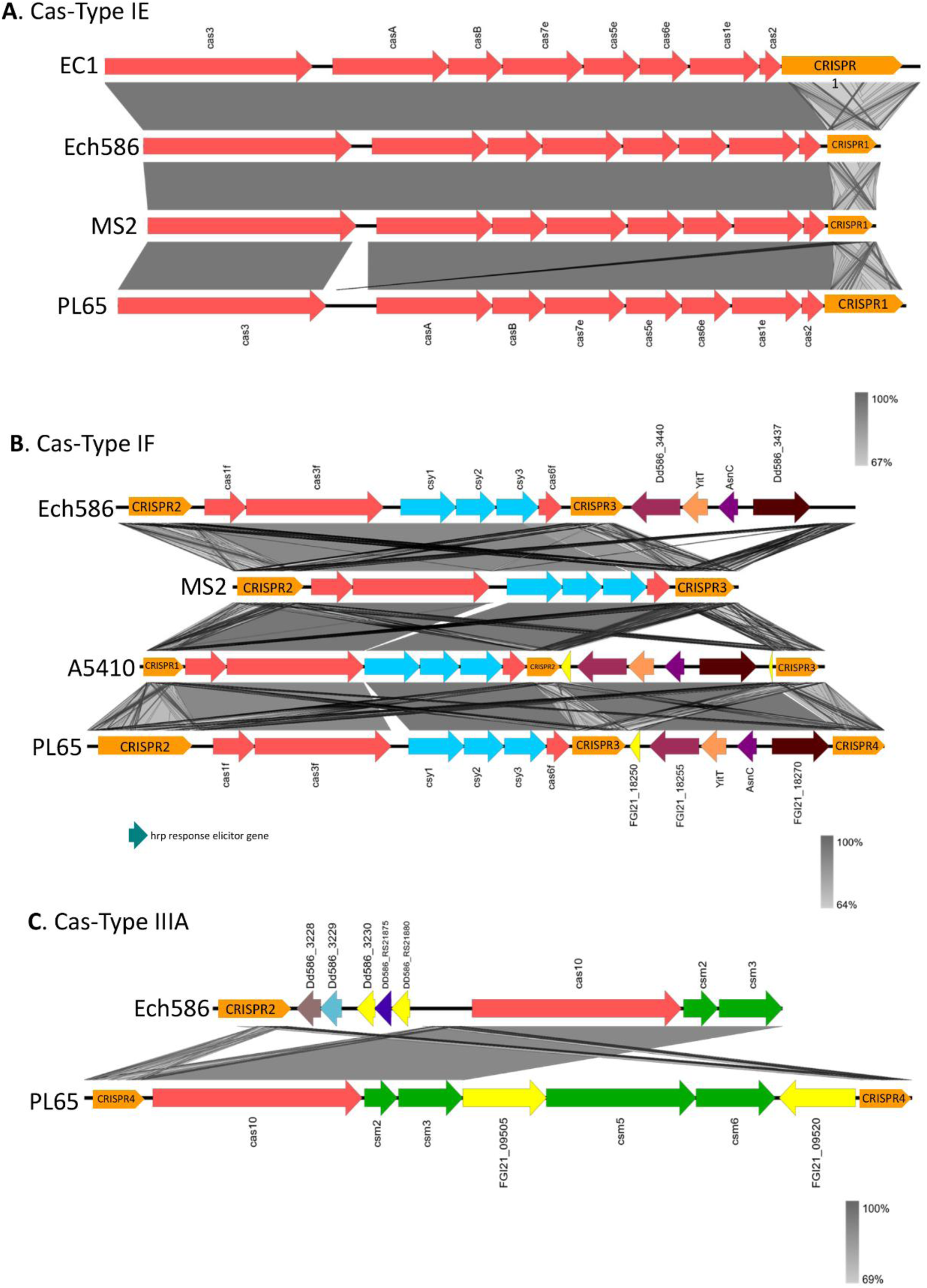
Diagram of the clustered regularly interspaced short palindromic repeats (CRISPR) with CRISPR associated proteins (Cas) system in five *Dickeya zeae* strains. (**A**) indicates the subtype I-E CRISPR-associated protein, (**B**) indicates the subtype I-F CRISPR-associated protein and Figure 14C indicates the type III-A CRISPR-associated protein. Orange arrows represents CRISPR repeats.

**Table 3.**
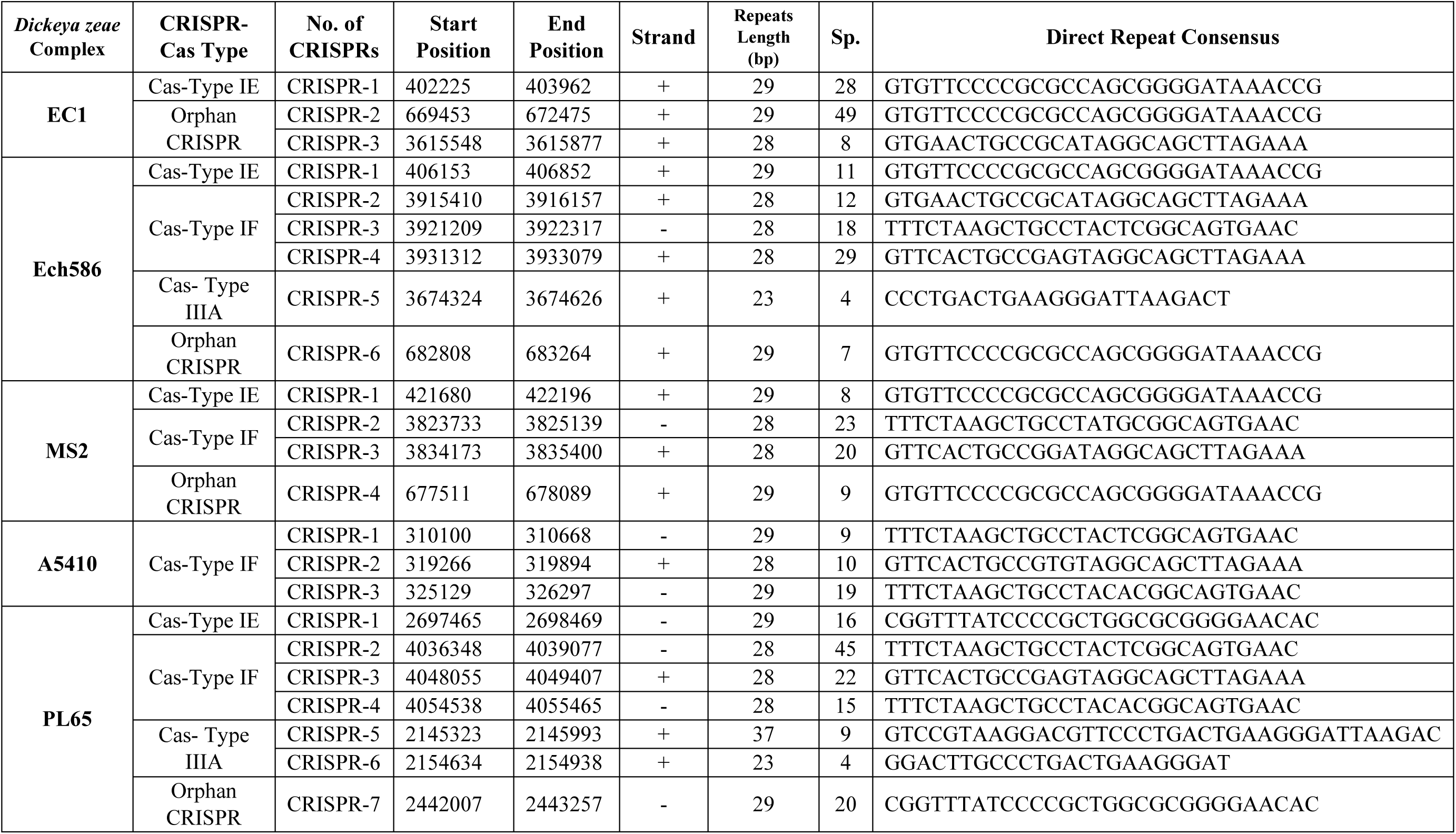
Detailed description of main features of CRISPR found in *Dickeya zeae* strains including EC1.

Interestingly, the A5410 genome harbored only the subtype I-F CRISPR-Cas. The type III-A CRISPR-Cas system was only found in PL65 and Ech586 genomes (**Figure 12**). While the genome of PL65 harbored two hypothetical proteins between the genes *cms*3 and *cms*5 and the other between *cas*6 and CRISPR region. PL65 and Ech586 genomes harbored the type III-A CRISPR-Cas system. Finally, we observed the presence of orphan CRISPRs that were distant from the *cas* operon regions—EC1 contained two orphan CRISPRs loci, whereas Ech586, MS2, and PL65 possessed one orphan CRISPR. The Orphan CRISPRs seem to operate their function far away from the *cas* locus, but they can be nonfunctional (Zhang et al. 2017).

The main characteristics of CRISPRs, such as position, length of direct repeats, number of spacers, and orientation, are provided in **Table 3**. Generally, the length of direct repeats within all CRISPRs were 28- to 29-bp, except for the direct repeats (DRs; 37 bp) of CRISPR located within type III-A CRISPR-Cas system of PL65. We observed the shortest CRIPSRs with a size of 302 bp — predicted for CRISPR5 in Ech586. The largest CRISPRs with a size of 3,022 bp—predicted for CRISPR2 in EC1. The highest number of 49 spacers were detected in the orphan CRISPR of EC1.

#### 3.7.8. Secondary metabolites within the *D. zeae* complex

The *D. zeae* complex revealed several genes involved in the synthesis of secondary antimicrobial-function metabolites. We used antiSMASH 4.0 server (Blin et al., 2017) to screen antimicrobials components. Four secondary metabolite biogenesis genes were predicted within *D. zeae* complex genomes and are summarized in **Table S3**. The five genomes harbor three well-known secondary metabolite biogenesis clusters [ind-vfm-expI, chrysobactin, and achromobactin], produced by *Dickeya* genus. The *ind-vfm-expI* genes are responsible for synthesis of the indigoidine molecule and the quorum-sensing mechanism (Charkowski et al., 2012; Nasser et al., 2013). The chrysobactin and achromobactin genes are involved in the biosynthesis of siderophores (Reverchon et al., 2013). Five strains also possess the gene cluster involved in the biosynthesis of cyanobactin-related molecules, which confers cytotoxicity. Further, seven clusters detected by AntiSMASH were found only in EC1, A5410, PL65, and Ech586; the bacteriocin synthesis cluster is present in all strains, except EC1. Beta-lactone containing protease inhibitor gene was predicted in all strains, except EC1. The arylpolyene biosynthesis cluster was identified in A5410, PL65, and Ech586. The bicornutin A1/A2 biosynthetic gene cluster [W909_RS19810, RS06850- RS06795], oocydin A biosynthetic gene cluster [W909_RS17185-17265], and the zeamine biosynthetic cluster [W909_RS19800, RS06540-RS06500] were found only in EC1 strains isolated from rice. The luminmide biosynthetic gene [FGI04_3605] was present only in A5410 strain isolated from pineapple.

### 3.7. Phenotypic comparison for *D. zeae* complex

We included three *D. zeae* strains [A5410, A5422 (NCPPB 2538=CFBP 2052), and PL65] for the phenotypic comparison assays. NCPPB 2538=CFBP 2052 was added as a reference. Firstly, we confirmed no difference among the three selected strains in the growth curve (**Figure S5**). We observed that A5410, A5422 (NCPPB 2538=CFBP 2052), and PL65 genomes harbored the pectate lyases (Pel cluster), cellulases (Cel5Z, CelH, and celY), and proteases (Prt cluster) genes. The selected three strains were able to produce protease, pectate lyases and cellulases assays on the plate assays (**Figure 13**). No significant difference was estimated (p>0.05) among the three strains in their ability to produce protease, pectate lyases and cellulases assays (**Figure 13**).

**Figure 13.**
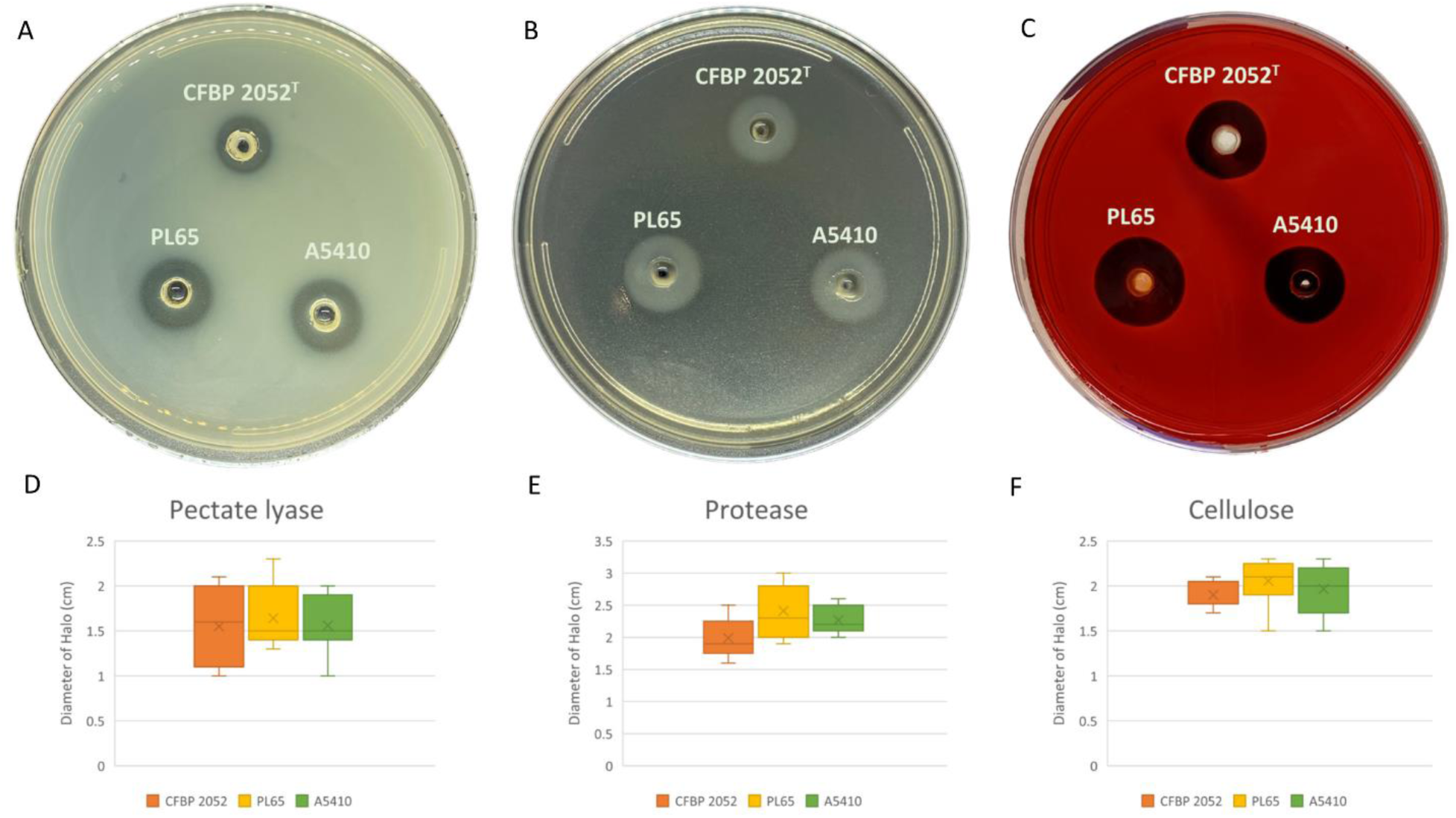
Extracellular cell wall degrading enzymes produced by three *D. zeae* strains. (**A)** The tree *D. zeae* strains on the pectate lyase assay plate; (**B)** The tree *D. zeae* strains on the protease assay plate; (**C)** The tree *D. zeae* strains on the cellulose assay plate. Samples of 50 µl of overnight culture were added to the assay plate wells (3 mm in diameter) and incubated at 28 °C. The Pel assay plates were treated with 4 N HCl after 10 h. The Cel assay plate was stained with 0.1% (w/v) Congo Red for 10 h and decolored with 5 M NaCl. The Prt assay plate was observed after 24 h without any further treatment. (**D)** Production of pectate lyase from *D. zeae* strains; **(E)** Production of protease from *D. zeae* strains; **(F)** Production of cellulose from *D. zeae* strains.

We observed that enterobacterial common antigen (ECA), capsular polysaccharide (cps), lipopolysaccharide, and extracellular polysaccharide (EPS) clusters were highly conserved in the three genomes. Moreover, the flagellar biosynthesis and chemotaxis proteins and Type IV pilus biogenesis proteins were found in all three genomes.

The biofilm formation, polysaccharide production, and mobility assays were performed. The selected strains were able to produce biofilm. There was no statistically significant difference for biofilm formation among three strains (p>0.05). Strain CFBP 2052 generated the highest exopolysaccharide on solidSOBG, while strain PL6 showed the smallest colony growth on solidSOBG. PL65 and A5410 generated nearly similar swimming and swarming zones, which were greater in comparison to CFBP 2052 (**Figure 14**). The results of EPS production assay and mobility assays showed a statistically significant difference with p<0.01.

**Figure 14.**
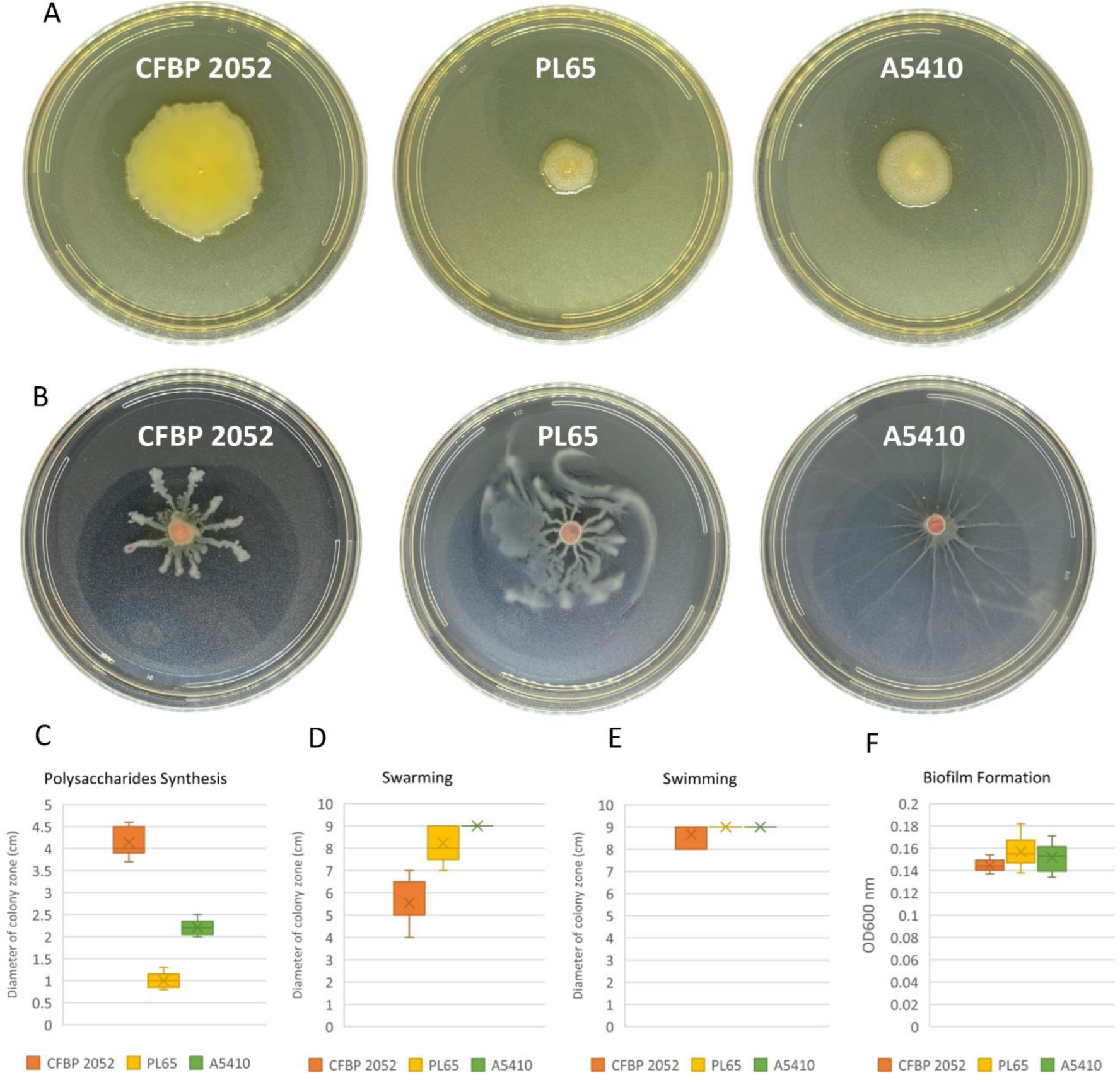
Exopolysaccharide production and motility of three *D. zeae* strains. (**A)** Exopolysaccharide (EPS) production of cells grown in SOB medium of three *D. zeae* strains and bacterial colony was spotted on the surface of SOB medium and the colony diameter measured 24 h later; (**B)** swarming capacity of three *D. zeae* strains was observed in semi solid medium, after 24 h at 28 °C; (**C**) production of exopolysaccharides from *D. zeae* strains; (**D)** capability of swarming of *D. zeae* strains; (**E)** capability of swimming of *D. zeae* strains; (**F**) ability of biofilm formation of *D. zeae* strains.

### 3.8. Pathogenicity assays

We evaluated pathogenicity on two different host plants, following the phenotypic characterization of three selected bacteria to understand the relationship between the phenotypic features and pathogenicity. We included (NCPPB 2538=CFBP 2052) type strain as a reference. We performed pathogenicity assays on taro corms and pineapple leaves based on the host of the strain PL65 and A5410. Results showed that PL65, A5410, and A5422 strains can infect all the tested host plants (**Figure 15**). Also, all three strains produced symptoms on the taro corm. The PL65 developed symptoms within just 6 hours after inoculation—faster than the other two. The results of the taro corm pathogenicity assay showed a statistically significant difference with p<0.01. PL65 strain, isolated from taro, produced greater tissue maceration compared to other two strains (**Figure 15**).

**Figure 15.**
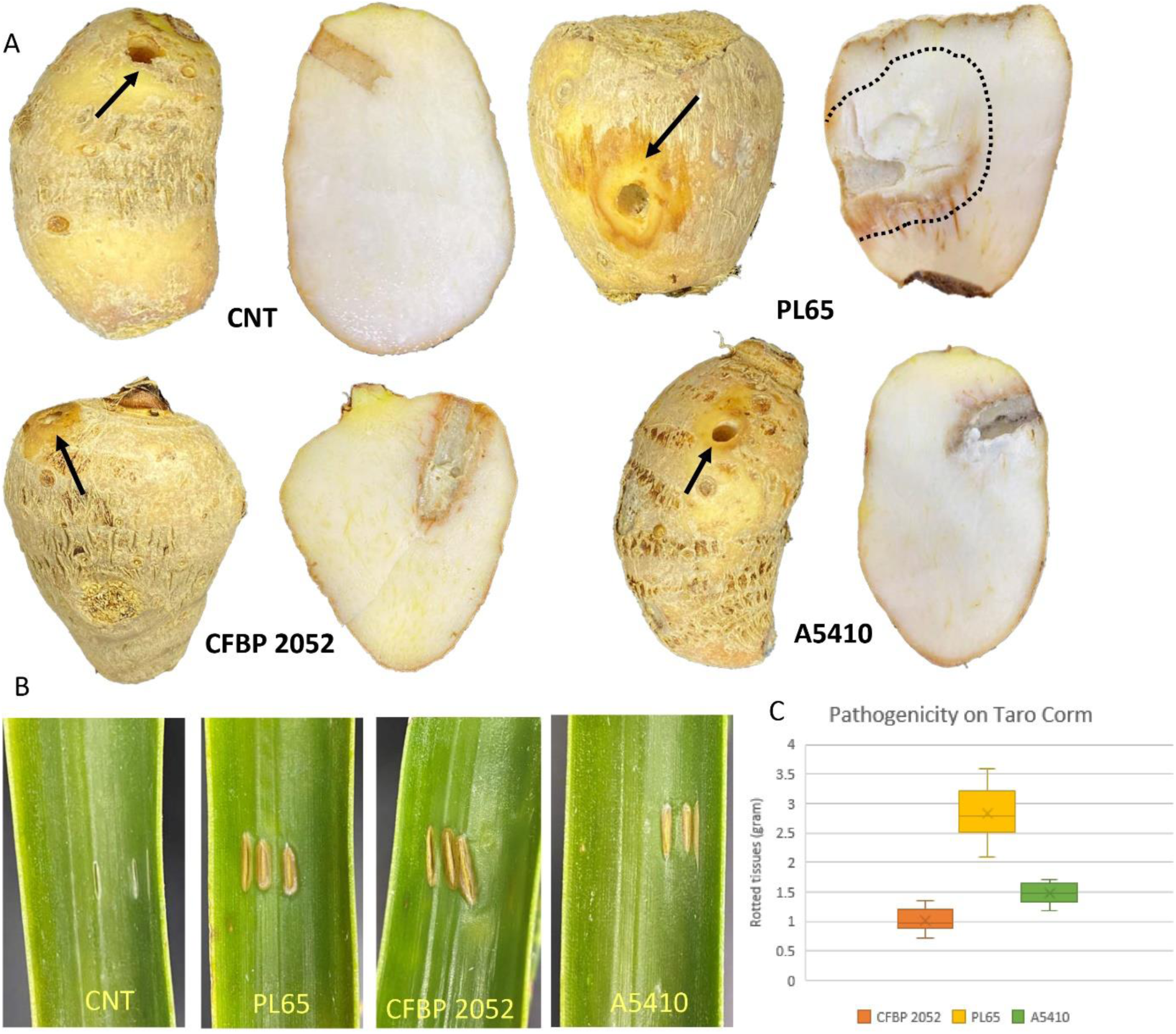
Pathogenicity of *D. zeae* strains on taro corm and pineapple leaf. (**A)** Picture of representative samples of infected plants and control plant (CNT) after incubation. Rotten area was indicated by black arrow and black cut line for taro corm. (**B)** Representative sample of infected plants and control plant (CNT) after incubation of pineapple leaf. **(C)** Quantification of soft rot mass on taro corm.

## 4. DISCUSSION

*Dickeya zeae* is a diverse and complex group within the genus *Dickeya* (Marrero et al 2008). In recent years, the taxonomic position of some strains of *Dickeya* has been changed. Recently, Wang et al. (2020) described a new species of *D. oryzae* in the *D. zeae* complex with a small margin of species demarcation threshold—the strains ZYY5, EC1, ZJU1202, DZ2Q, NCPPB 3531, and CSL RW192 were separated from *D. zeae* (Wang et al. 2020). Recently, a novel strain of *D. zeae* was isolated from rice showing distinct characteristics (Zhang et al, 2020). Based on our research outcomes, we observed that *D. zeae* still comprising the novel species but with little species threshold margins (Boluk and Arif, unpublished information).

One of the main virulence factors for soft rot bacteria is extracellular enzymes (Hugouvieux-Cotte-Pattat et al., 1996). The PCWDEs (pectinases, cellulases, and proteinases) are considered the essential virulence factors for the host colonization and disease development (Charkowski et al., 2012; Davidsson et al., 2013). Pectinase enzymes such as pectate lyases (*Pel*), pectin methylesterases (*Pem*) and pectin lyases (*Pnl*) have been studied on *D. dadantii* 3937 (Hugouvieux-Cotte-Pattat et al., 1996). Pectinase enzymes play a significant role in the virulence and tissue maceration (Hugouvieux-Cotte-Pattat et al., 1996). Previous comparative genomic analyses revealed that the genes related to the production of PCWDEs include multiple pectate lyases (*Pel*ABCDEILNWXZ and *Pel*10), pectin lyases (*Pnl*GH), polygalacturonases (*Peh*KNVWX), pectin methyl esterases (*Pem*AB), pectin acetyl esterases (*Pae*XY), feruloyl esterases (*Fae*DT), rhamnogalacturonate lyases (*Rhi*EF), and one periplasmic endogalactanase (*Gan*A) exist in various *Dickeya* species and are highly conserved (Zhou et al., 2015; Duprey et al., 2019)—our analyses demonstrated concordant results.

Gram-negative bacteria have evolved several complex secretion systems to translocate a wide range of extracellular enzymes and effector proteins from the periplasm across the outer membrane (Costa et al., 2015). The structural and mechanistic features of the types I–VI were described in the gram-negative bacteria (Costa et al., 2015). The proteases, which is another crucial virulence, are secreted by the T1SS known as the *prt*DEF operon (Toth et al., 2006; Charkowski et al., 2012). Zhou et al. (2015) reported that *D. paradisiaca* Ech703 did not harbor the *prt*DEF genes among the *Dickeya* species that is concordant to our findings. Previous studies have revealed that many Gram-negative bacteria use the type II secretion system (T2SS) to translocate extracellular proteins such as pectinase and cellulase enzymes (Filloux 2004; Korotkov et al., 2011). T2SS gene cluster (*out*SBCDEFGHIJKLMO) is well-conserved among *Dickeya* species (Zhou et al., 2015; Duprey et al., 2019)—similar results were obtained in our analyses with all five strains isolated from diverse hosts including rice, banana, pineapple, taro, and Philodendron (**Figure S3**). The T3SS plays a vital role in modulating plant defenses for several plant bacterial pathogens, including *Pseudomonas syringae*, *Erwinia,* and *Xanthomonas* sp., (Deslandes and Rivas 2012). However, recent studies indicated that a few *Pectobacterium* species such as *P. parmentieri, P. wasabiae* and *D. paradisiaca* lack the T3SS cluster (Kim et al., 2009; Nykyri et al., 2012; Arizala and Arif, 2019; Duprey et al., 2019), and stated that the T3SS is not necessary for *Pectobacterium* species for disease development (Arizala and Arif, 2019). We found that T3SS cluster was present in all strains—EC1, MS2, Ech586, A5410—except PL65 isolated from taro (**Figure 10**). The role of the *vir*B (T4SS) operon was demonstrated in *P. atrosepticum* as a virulence factor (Bell et al., 2004). *Dickeya dadantii* 3937 and *D. fangzhongdai* (ND14b, M074, and M005) encodes both types of T4SS, a *Vir*D2/*Vir*D4/*Trb* locus and *vir*B operon (Pédron et al., 2014). *Dickeya zeae* complex species possessed only one type of T4SS, *vir*B operon. Interestingly, PL65 and MS2 harbored *vir*B1 and *vir*B2 (**Figure 11**). Previous studies indicated that *vir*B1 forms a borehole in the peptidoglycan layer that enables complex T4SS assembly to occur, and the proteins *Vir*B2 and *Vir*B5 constitute T4SS extracellular pilus (Fronzes et al., 2009; Chandran et al., 2009).

Flagellar biosynthesis and chemotaxis proteins were found in all five strains (EC1, Ech586, MS2, A5410, and PL65). Previously, Zhou et al. (2015), have proved that EC1, DZZ2Q, and ZJU1202 strains isolated from rice possessed the flagellar biosynthesis genes clusters. In many plant-pathogenic bacteria, flagella proteins are responsible for cell motility and secretion and vesicular transport (Jahn et al., 2008), and the motility lends to virulence (Chesnokova et al. 1997; Mulholland et al. 1993; Panopoulos and Schroth 1974; Tans-Kersten et al. 2001). Flagella are used for both swimming and swarming motility (Yi et al. 2000). Individual swimming cells perceive a chemical signal via methyl-accepting chemotaxis proteins responsible for cell motility and signal transduction. (Yi et al., 2000). It has been demonstrated that the mutation within chemotactic genes (*cheW, cheB, cheY*, and *cheZ*) caused a substantial reduction in swimming motility (Antúnez-Lamas et al., 2009). The PL65 and A5410 isolated from taro and pineapple, respectively, showed similar swimming ability, but PL65 caused the highest virulence against taro corm among the three strains tested (**Figure 14-15**). Jahn et al. (2008) have proved that the mutation of the *fli*A gene encoding a sigma factor obstructed the bacterial motility and limited *Pel*s production and the bacterial attachment to plant tissues in *D. dadantii* 3937. These results show that flagellar biosynthesis and chemotaxis proteins are associated with virulence. Another virulence factor studied in plant-pathogen bacteria, such as *Ralstonia* and *Xylella*, is type IV pilus (Burdman et al., 2011). The type IV pilus assembly encoded by *pil* genes is responsible for twitching motility in the *Pseudomonas aeruginosa* and *D. aquatica* (Maier and Wong, 2015; Duprey et al., 2019). We found this *pil* gene cluster in all *D. zeae* strains.

*Dickeya* species produce secondary metabolites such as the thiopeptide, cyanobactin, zeamine and oocydin (Zhou et al., 2011; Matilla et al., 2012; Zhou et al., 2015; Alic et al., 2019; Duprey et al., 2019). The PKs (polyketides) and NRPs (nonribosomal peptidesres) are two representative classes of enzymes that synthesize essential secondary metabolites (Blin et al., 2013). The zeamine gene cluster is well known among the secondary bioactive metabolites for *Dickeya* species such as *D. fangzhongdai, D. solani,* and *D. oryzae* (previously known as *D. zeae*) (Zhou et al., 2015; Zhang et al., 2018; Alic et al., 2019; Duprey et al., 2019). *Dickeya oryzae* (ZJU1202, DZ2Q, EC1, and ZYY5) strains isolated from rice possessed the zeamine gene cluster among the *D. zeae* complex strains (Zhou et al., 2015, Wang et al. 2020). This distinctive difference among the *D. zeae* complex strains was used to define the novel species of *Dickeya* (Wang et al. 2020). Additionally, we observed that strain EC1 produced the antifungal compound oocydin via non-ribosomal peptide synthases (NRPS) and polyketide synthases (PKS). Oocydin is responsible for the strong antimicrobial activity against plant pathogenic fungi and oomycetes (Matilla et al., 2012). Similar cluster sequences were also present in other *Dickeya* species such as *D. solani*, *D. dianthicola*, *D. zeae*, *D. chrysanthemi*, *D. fangzhongdai* and *D. paradisiaca* (Duprey et al., 2019; Alic et al., 2019). The antioxidant indigoidine is a well-known secondary metabolite, produced by all *Dickeya* (Nasser et al., 2013). Moreover, we predicted that the *D. zeae* complex harbored the antioxidant indigoidine.

The polysaccharides synthesis clusters such as capsular polysaccharide, extracellular polysaccharide and lipo-oligo/polysaccharide have been described as one of the important virulence factors that allows bacteria to bind to the host cell surface (Kuhn et al., 1988; Reeves et al., 1996; Roberts, 1996; Bell et al., 2004; Panda, 2006; Toth et al., 2003; Toth et al., 2006; Nykyri et al., 2012; Arizala and Arif, 2019). The EPS is a main component for the bacterial biofilm matrix and responsible for adhesion to the plant surfaces (Flemming and Wingender, 2010). *Pectobacterium* genus harbored the EPS biosynthesis cluster (*wza-wzb-wzc-wzx*) (Arizala and Arif, 2019). In this study, we found that all five strains harbored the EPS cluster, besides, we did not observe any difference in biofilm formation among these strains. The *cps* (capsular polysaccharide) cluster which is not crucial for living was not observed in some *Pectobacterium* members (Arizala and Arif, 2019). We determined that strain EC1 possessed the LOP and EPS genes cluster; however, the *cps* cluster was absent. Moreover, in the SOBG medium assay, the EC1 had the highest ability to produce polysaccharides. There are extra factors involved in biofilm formation such as mobility (swimming and swarming) and twitching motility.

In this study, some important genes were also identified and predicted to play functional roles. These genes were annotated and associated with the functions of production of antimicrobials, nitrogen fixation, and the uptake and catabolism of aromatic compounds. We demonstrated that only A5410 isolated from pineapple harbored the nitrogen fixation cluster (*nif*ABCDEHKLMNSTUVQWXYZ), the *ars*C (arsenic resistant gene) gene and carboxymuconolactone decarboxylase gene. The nitrogen fixation cluster was also present in *D. solani* and *P. atrosepticum* (Bell et al., 2004; Tsror et al., 2012; Golanowska, 2015). Carboxymuconolactone decarboxylase participates to catalyze aromatic compounds to produce acetyl- or succinyl-CoA, in prokaryotes and yeast (Cheng et al., 2017). Carboxymuconolactone decarboxylase was present in *Azotobacter vinelandii, Acinetobacter calcoaceticus* and *Pseudomonas putida* (Yeh et al., 1981). Bacterial survival within a specific environment is linked to how bacteria cope with toxic compounds. Acquisition of arsenic clusters (*ars*) confer the ability of the bacteria to resist against the concentrations of inorganic arsenic present in the environment (Fekih et al., 2018). A recent study showed that *P. atrosepticum, P. brasiliense*, *P. peruviense* and *P. versatile* (formerly proposed as Candidatus *Pectobacterium maceratum*) present four genes of arsenic clusters: *arsC*, *arsB*, *arsR* and *arsH* (Bell et al., 2004; Arizala and Arif, 2019). The *arsC* and *arsH* genes in the *Pectobacterium* are vital for living in the arsenic rich ecology (Arizala and Arif, 2019).

The CRISPR-Cas (Clustered Regularly Interspaced Short Palindromic Repeats and the CRISPR-associated proteins) is widely distributed and found in at least half of the bacteria and almost all archaea (Haft et al., 2005). The CRISPR-Cas is defined as the immune system that can protect against bacteriophages and foreign plasmids (Mojica et al., 2005; Makarova et al., 2011; Rath et al., 2015). Three types of CRISPR-Cas systems (type I, II, and III) were classified based on the *Cas*3, *Cas*9, and *Cas*10 proteins. The type I comprises further subtypes (e.g., I-A to I-F), each is characterized by a specific set of proteins (Przybilski et al., 2011). Two types of CRISPR-Cas systems (cas-Type III and cas-subtype IE/IF) were described among *D. zeae* complex species. Previously, the cas-subtype IE was observed in *D. fangzhongdai, D. dadantii*, *D. zeae*, and *D. paradisiaca,* and the cas-subtype IC was identified in *D. dadantii* and *D. dianthicola*. In our analyses, orphans CRISPRs were found throughout the 4 analyzed *Dickeya* genomes (**Table 3**). The orphan CRISPRs described that some CRISPRs seem to exert their function far away from the *cas* locus, although they might not be functional (Zhang et al., 2017). Similarly, orphan CRISPRs were also found in some *Pectobacterium* species and these might be functional or just remnants of previous CRISPR-Cas systems (Arizala and Arif, 2019).

As a conclusion, in this study, we present two high quality complete genome sequences of novel *D. zeae* strains PL65 and A5410 isolated from taro and pineapple in Hawaii. The detailed comparative genomic analyses were performed by using the aforementioned strains along with three other strains retrieved from the NCBI GenBank genome database. For taxonomic and phylogenomic analyses, representative strains from other species were also included. Several groups of virulence genes, such as the cell wall-degrading extracellular enzymes genes, T1SS gene cluster *prtDEF*, T2SS gene cluster (*out* gene cluster), T5SS gene cluster, T6SS gene cluster, flagella and chemotaxis gene clusters,some polysaccharides synthesis clusters, and type IV pilus gene cluster are highly or fully conserved in all five genomes isolated from different hosts. Interestingly, T3SS and T4SS gene clusters were absent in the strain PL65 isolated from taro. Additionally, we also found that the T4SS gene cluster was absent in MS2. Importantly, a range of unique genes associated with virulence were identified in A5410, such as arsenic gene, nitrogen fixation cluster. Intriguingly, the *zms* (zeamine) genes cluster and oocydin genes cluster were found only in EC1 which was isolated from rice.

## ACKNOWLEDGMENTS

This work was supported by the USDA National Institute of Food and Agriculture, Hatch project 9038H, managed by the College of Tropical Agriculture and Human Resources. Research was also supported by NIGMS of the National Institutes of Health under award number P20GM125508. Strain A5410 was obtained from Pacific Bacterial Culture Collection that is maintained by the National Science Foundation (Award No. 1561663). The content is solely the responsibility of the authors and does not necessarily represent the official views of the funding agencies.

## SUPPLEMENTAL MATERIALS

**Figure S1.** Pairwise heatmap based upon the average nucleotide identity (ANI) and digital DNA-DNA hybridization (dDDH) values in terms of percentage among the fourteen *Dickeya* species and a *Pectobacterium* species. The upper diagonal displays ANI data whereas the lower diagonal depicts the *in silico* DDH data. Cut-off values for species delineation are 95-96 and 70 % for ANI and dDDH, respectively.

**Figure S2. (A**). The dDDH phylogenetic tree were inferred with FastME 2.1.6.1 from TYGS_GBDP distances calculated from genome sequences (Kolthoff and Goker, 2019). (**B**). The ANI phylogenetic tree was generated for the *Dickeya* species strains based on whole genome alignment using the neighbor–joining method. The Jukes–Cantor model was used for analysis with 1,000 bootstraps.

**Figure S3.** Comparison of the genetic organization of type I secretion system (**A**) and type II secretion system (**B**) among five *Dickeya zeae* strains. The arrow position represented forward/reverse gene orientation. Arrow color signified specific gene composition within the T1SS and T2SS. Gene names were provided at the top and bottom of the linear graph. A pairwise alignment between the linear sequences was rendered based upon BLAST algorithm with cut-off values from 66% to 100% and 82% to 100%, T1SS and T2SS, respectively. Regions with higher nucleotide identity were displayed with a shaded grey.

**Figure S4.** Comparison of the genetic organization of type V secretion system (**A**) and type VI secretion system (**B**) among five *Dickeya zeae* strains. The arrow position represented forward/reverse gene orientation. Arrow color signified specific gene composition within the T5SS and T6SS. Gene names were provided at the top and bottom of the linear graph. A pairwise alignment between the linear sequences was rendered based upon BLAST algorithm with cut-off values from 68% to 100%. Regions with higher nucleotide identity were displayed with a shaded grey.

**Figure S5.** Bacterial growth curve of *Dickeya zeae* strains (CFBP 2052, A5410 and PL65). Bacterial cultures were grown with Nutrient Broth at 37°C, with shaking. These data represent three separate experiments.

**Table S1.** Eighty-six virulence-related genes (cell wall-degrading enzyme genes) from *Dickeya* sp. and their locus tag within each genome.

**Table S2.** Flagellar and chemotaxis genes, twitching motility genes, polysaccharide biosynthesize gene clusters from *Dickeya* sp. and their locus tag within each genome.

**Table S3.** The secondary metabolite gene clusters identified with AntiSMASH in five genomes of *Dickeya* sp.

