## Supplementary figures and images for "Genomic and phenotypic biology of novel strains of *Dickeya zeae* isolated from pineapple and taro in Hawaii: insights into genome plasticity, pathogenicity, and virulence determinants"

### Figure S1

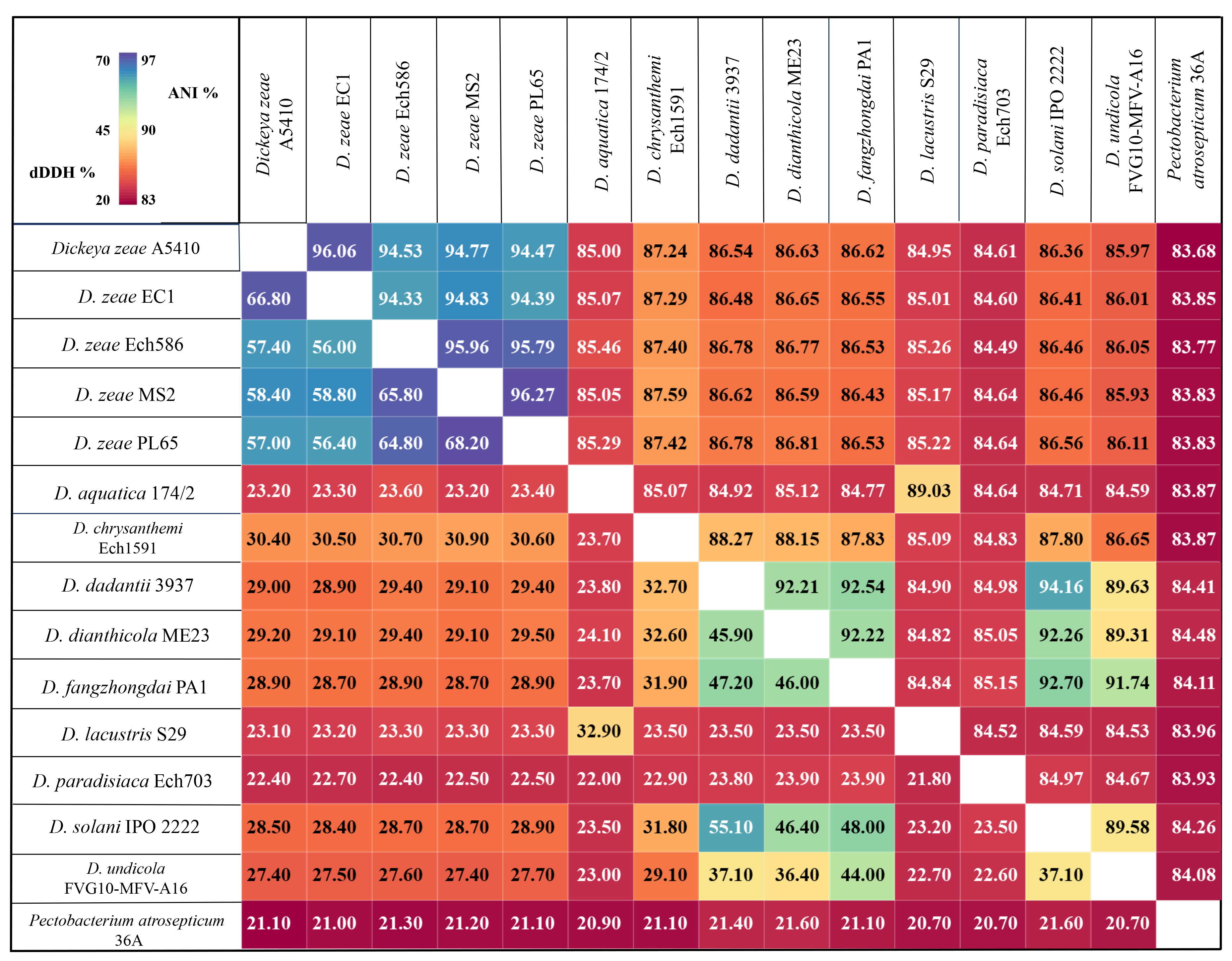

### Figure S2

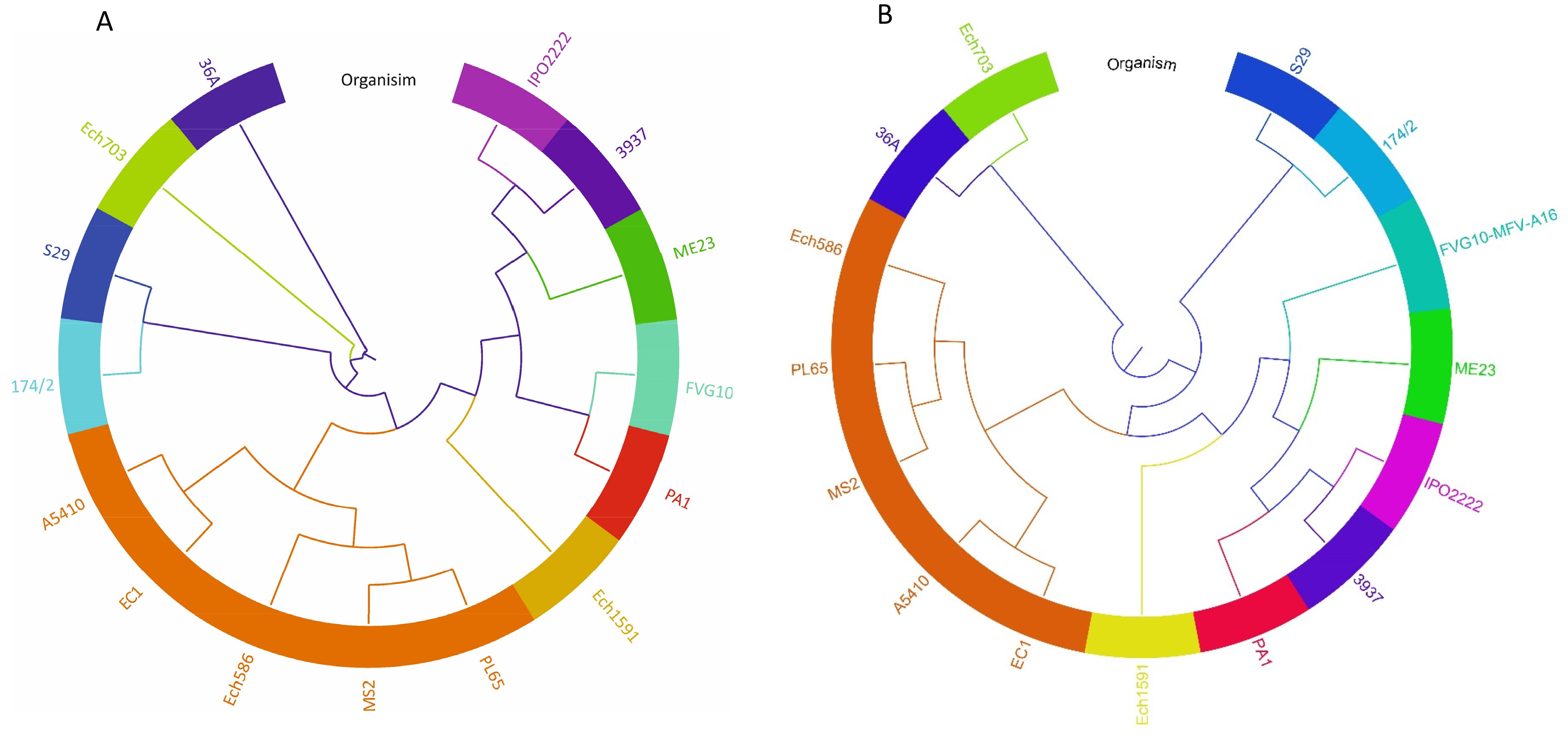

### Figure S3

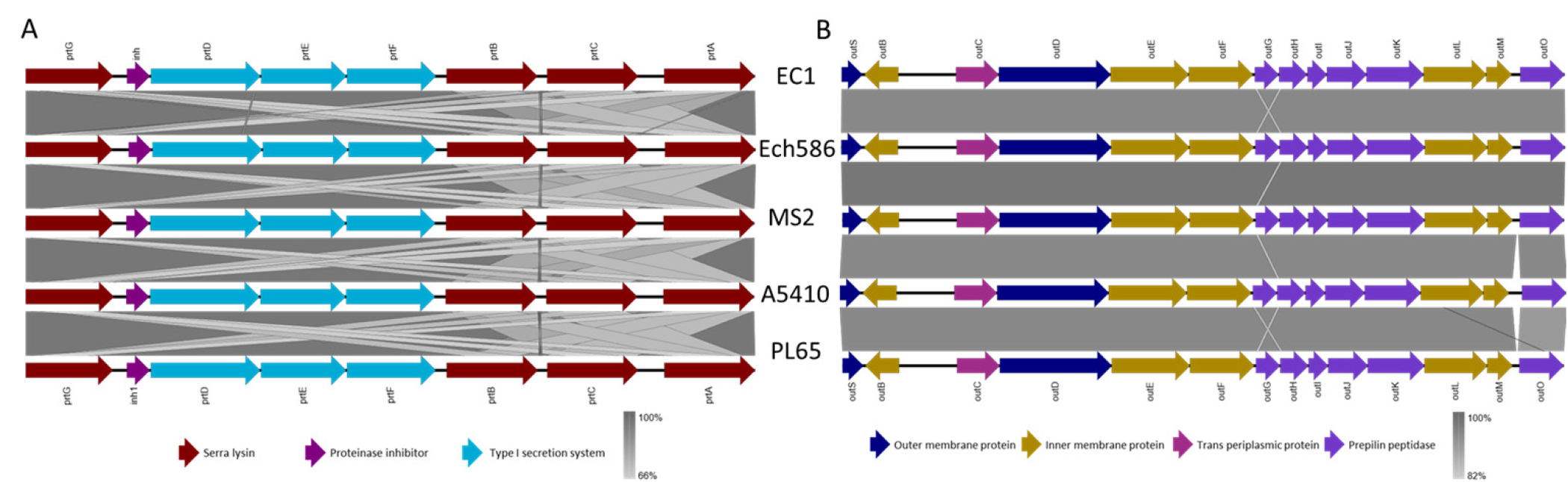

### Figure S4

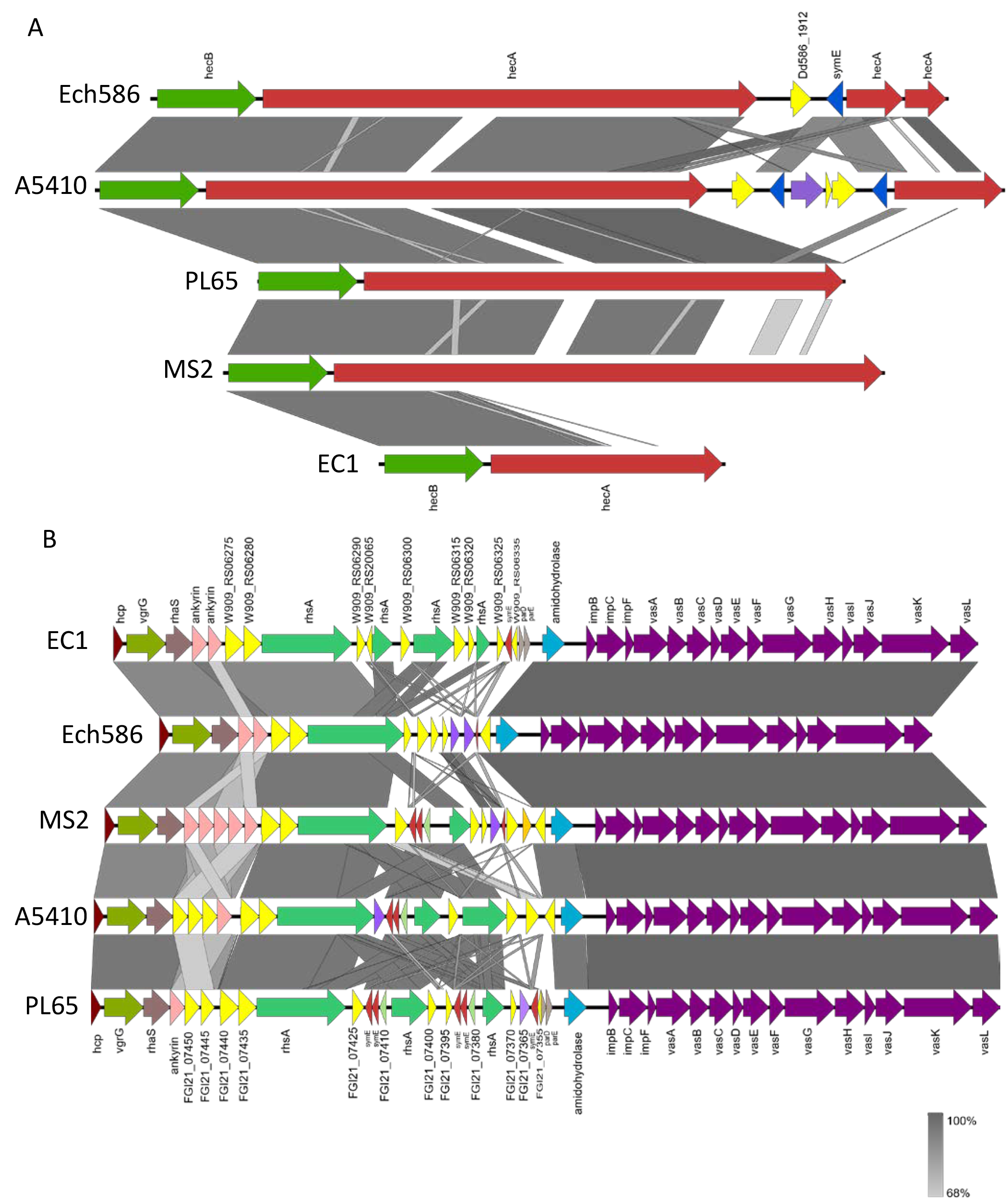

### Figure S5

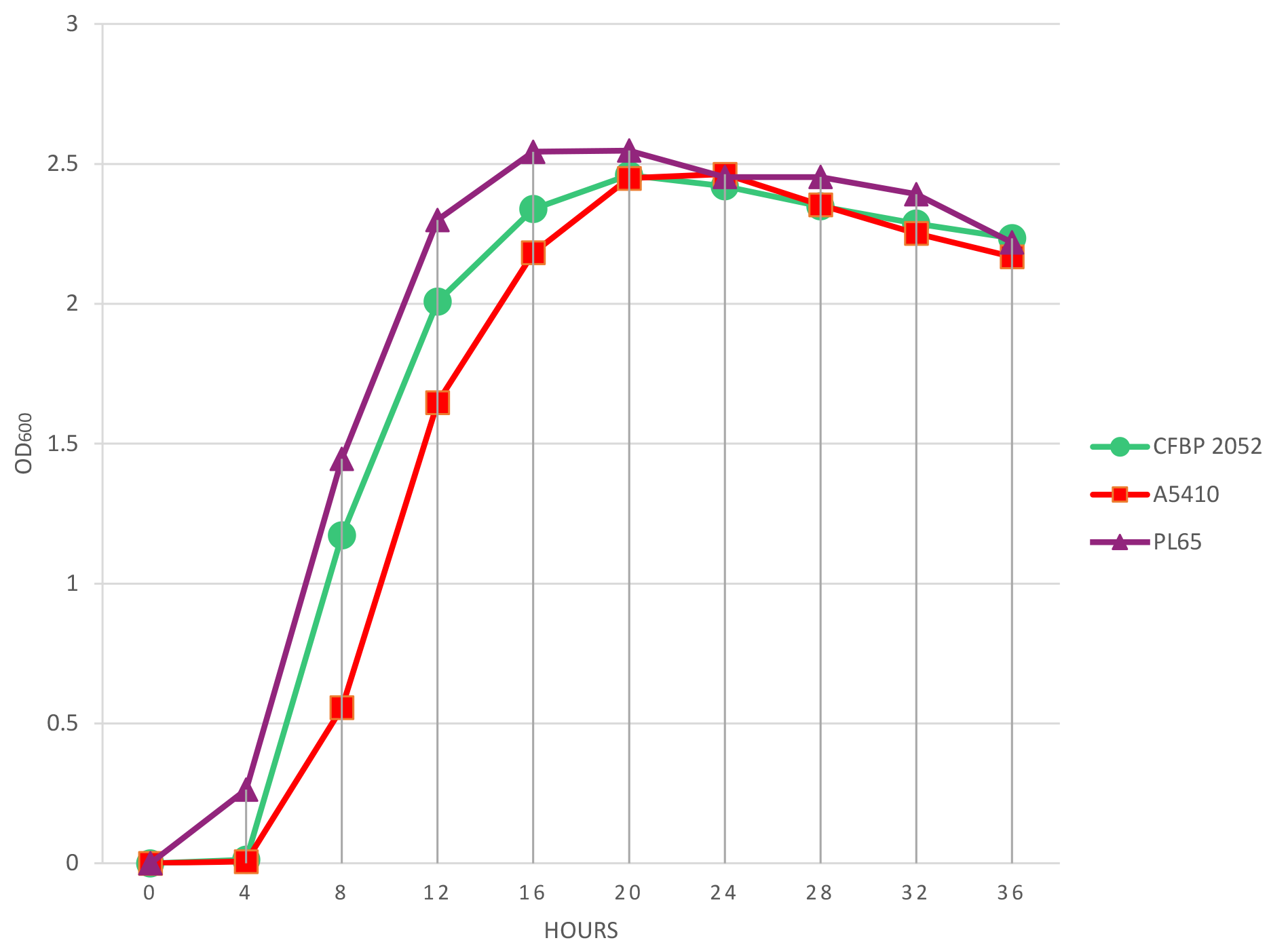
