## Supplementary material for "Genomic and phenotypic biology of novel strains of *Dickeya zeae* isolated from pineapple and taro in Hawaii: insights into genome plasticity, pathogenicity, and virulence determinants": Table S3

| **Secondary metabolite gene clusters** | A5410 | PL65 | EC1 | Ech586 | MS2 |
| --- | --- | --- | --- | --- | --- |
| Ind-vfm-expI | + | + | + | + | + |
| Achromobactin | + | + | + | + | + |
| Chrysobactin | + | + | + | + | + |
| Cyanobactin | + | + | + | + | + |
| Betalactone | **+** | + | - | + | + |
| Bacteriocin (TfuA) | + | + | - | + | + |
| Arylpolyene | + | + | - | + | - |
| Bicornutin A1/A2 | - | - | + | - | - |
| Oocydin | - | - | + | - | - |
| Zeaemine | - | - | + | - | - |
| Luminmide | + | - | - | - | - |

**Table S3**. The secondary metabolite gene clusters identified with AntiSMASH in five genomes of *Dickeya* sp.
